# CCDC81 is a conserved microtubule-binding protein that maintains centriole integrity

**DOI:** 10.64898/2026.11.16.766940

**Authors:** Umut Batman, Daria Khuntsariya, Susanne Borgers, Paul Guichard, Virginie Hamel

**Affiliations:** University of Geneva, Department of Molecular and Cellular Biology, Faculty of Sciences, Geneva, Switzerland

**Keywords:** A-C linker, centriole, basal bodies, *Chlamydomonas*, human cells, ultrastructure expansion microscopy, iU-ExM

## Abstract

Centrioles are microtubule-based organelles that exhibit a highly conserved architecture that is essential for their function. However, the molecular mechanisms maintaining their structural integrity remain incompletely understood. Here, we characterize CCDC81 as a conserved microtubule-binding protein that contributes to centriole organization. Using expansion microscopy, we localize CCDC81 to the A-C linker of *Chlamydomonas* basal bodies, where it associates with POC11, the homolog of human A-C linker protein CCDC77. We demonstrate that CCDC81 directly binds microtubules through its N-terminal HU domains, promotes microtubule bundling, and stabilizes microtubules by reducing depolymerization and increasing rescue frequency. Cryo-electron microscopy further reveals its association with the outer microtubule lattice. We identify distinct domains mediating CCDC81 interaction with POC11 and propose a molecular organization of this complex within the A-C linker. Importantly, CCDC81 also localizes to human centrioles, where its depletion disrupts centriole architecture and A-C linker organization, with defects exacerbated by co-depletion of CCDC77. Together, our findings establish CCDC81 as a conserved regulator of microtubule organization and centriole structural integrity.

## Introduction

Centrioles and basal bodies are evolutionarily conserved microtubule-based organelles that play essential roles in centrosome organization and in the formation of cilia and flagella (LeGuennec *et al*., 2021). In human cells, centrioles are barrel-shaped structures approximately 450 nm long and 250 nm in diameter (LeGuennec *et al*., 2021; Laporte *et al*., 2024), characterized by a ninefold radial symmetry of microtubule triplets. Each triplet consists of a complete A-tubule associated with two incomplete B- and C-tubules (Li *et al*., 2012; Klena *et al*., 2020). The remarkable stability of this architecture relies on a network of microtubule-associated proteins and structural complexes that maintain triplet cohesion and reinforce the centriolar wall (Le Guennec *et al*., 2020; Steib *et al*., 2020). Along their longitudinal axis, centrioles exhibit a polarized organization comprising proximal, central, and distal regions, each containing distinct molecular assemblies that contribute to centriole assembly, stability, and function (Laporte *et al*., 2024).

At the proximal end, the cartwheel, primarily organized around the ninefold symmetric SAS-6 scaffold, establishes centriole symmetry and provides the structural framework for procentriole assembly (Kitagawa *et al*., 2011; Van Breugel *et al*., 2011; Guichard *et al*., 2017). The cartwheel is connected to the microtubule triplets through the pinhead, while adjacent microtubule triplets are interconnected by the A-C linker, which extends along approximately 40% of the centriole length (Laporte *et al*., 2024; Bournonville *et al*., 2025). Cryo-electron tomography (cryo-ET) studies in evolutionarily distant organisms, including *Trichonympha*, *Paramecium*, and *Chlamydomonas*, have revealed a broadly conserved organization of the A-C linker, although its detailed architecture varies considerably between species (Li *et al*., 2012; Guichard *et al*., 2013; S. Li *et al*., 2019; Klena *et al*., 2020);. In human cells, Ultrastructure Expansion Microscopy (U-ExM) identified CCDC77, WDR67/TBC1D31, and MIIP as components of an A-C linker-associated complex, whose disruption compromises microtubule triplet cohesion, centriole integrity, and duplication (Laporte *et al*., 2024; Bournonville *et al*., 2025) More recently, by combining cryo-ET and single-particle cryo-EM of isolated *Tetrahymena thermophila* centrioles, Cai *et al*., (2025) identified two distinct A-C linker configurations along the centriolar microtubule triplets. In the proximal region, the A-link contains WRAP73 and CEP135, whereas the C-link comprises POC1 and WDR67/TBC1D31, with the latter forming a central filament connecting the two links. In the adjacent transitional region, which lacks both the cartwheel and inner scaffold, the A-link is largely preserved, while the C-link adopts a distinct organization involving unassigned β-propeller domains and a C-microtubule-binding domain. These findings reveal that the A-C linker is not a uniform structural bridge but rather a molecularly heterogeneous assembly whose composition and organization vary along the centriole. Importantly, several structural densities remain unassigned, indicating that the molecular composition of the A-C linker is still incomplete and that additional proteins likely contribute to its organization and interaction with centriolar microtubules.

Beyond the proximal region, centriole stability is further supported by the inner scaffold, a large structural assembly lining the luminal surface of the microtubule wall. Initially visualized by cryo-ET as a periodic, lattice-like structure extending along the central region of centrioles, the inner scaffold has subsequently been characterized through the combination of U-ExM and structural approaches. These studies have identified several associated proteins, including POC5, FAM161A, POC1A/B, centrins, CCDC15, and, more recently, AKIP1 (Le Guennec *et al*., 2020; Steib *et al*., 2020; Arslanhan *et al*., 2023; Sala *et al*., 2024; Sanada *et al*., 2026). Unlike the A-C linker, which connects neighboring microtubule triplets in the proximal region, the inner scaffold provides an internal structural framework that reinforces the centriolar microtubule wall. Consistent with this function, depletion of individual inner scaffold components can disrupt scaffold organization, compromise microtubule cohesion, and ultimately lead to centriole fracture (Steib *et al*., 2020; Arslanhan *et al*., 2023; Sala *et al*., 2024; Sanada *et al*., 2026). At the distal end, a luminal distal ring containing C2CD3, SSNA1, SFI1, and CEP135 contributes to distal centriole organization (Laporte *et al*., 2022, 2024; Bertiaux *et al*., 2025; Huang *et al*., 2025), while more peripheral proteins, including MNR, CEP90, and OFD1, participate in establishing the architecture required for distal appendage formation (Wang *et al*., 2018; Kumar *et al*., 2021; Le Borgne *et al*., 2022). Together, these findings reveal that centriole integrity relies on multiple spatially organized structural modules that collectively stabilize the microtubule barrel and support its functional specialization.

Despite these advances in resolving centriole architecture, the molecular composition of several structural assemblies remains incomplete, and the mechanisms by which individual proteins interact with and stabilize centriolar microtubules are still poorly understood. This is particularly relevant for the A-C linker, where distinct molecular configurations suggest that multiple proteins cooperate to maintain microtubule triplet cohesion. One potential contributor to this architecture is CCDC81, an evolutionarily conserved but poorly characterized centrosomal protein. Initially identified through centrosomal proteomic analyses, CCDC81 was subsequently shown to localize to centrosomes and basal bodies in human cells (Firat-Karalar *et al*., 2014; Liu, Liu and Nai, 2026). Its centrosomal targeting was mapped to a central region of the protein, while its depletion was reported to reduce both the frequency and length of primary cilia, suggesting a role in ciliogenesis (Liu, Liu and Nai, 2026). Interestingly, proteomic analysis of isolated *Chlamydomonas reinhardtii* basal bodies identified the CCDC81 homolog as the first most abundant protein (Hamel *et al*., 2017), suggesting that it may represent a major component of this organelle. Nevertheless, its precise localization within the centriole, its molecular properties, and its contribution to centriole architecture remain unknown.

CCDC81 belongs to an ancient family of proteins characterized by two tandem N-terminal domains related to the bacterial HU/IHF superfamily (Burroughs *et al*., 2017). In bacteria, HU proteins are well known for their ability to bind and organize DNA, contributing to chromosome compaction and nucleoid architecture. Phylogenetic analyses suggest that the eukaryotic HU domain family originated through an ancient lateral gene transfer event and was subsequently retained across diverse ciliated lineages, including Giardia, kinetoplastids, green algae, and animals, while being largely absent from organisms that have lost centrioles and cilia (Burroughs *et al*., 2017). Notably, the residues involved in DNA binding in bacterial HU proteins are not conserved in CCDC81, raising the possibility that these domains have acquired a distinct molecular function during evolution. The strong evolutionary association of this protein family with ciliated organisms suggests that such a function may be linked to microtubule-based structures. Consistent with this possibility, several members of the CCDC81-related protein family have been identified in association with axonemal microtubules. In *Tetrahymena*, a protein containing a CCDC81-related domain was identified within a CEP104-associated complex at the ciliary tip (Legal *et al*., 2023). Other structurally related proteins, including IJ34 and CCDC81B, have been associated with axonemal microtubule doublets, with IJ34 contributing to the organization of the inner junction between the A- and B-tubules (Kubo *et al*., 2023; Gao *et al*., 2024; Li *et al*., 2025). Furthermore, CCDC81 itself has been identified among microtubule inner proteins in sea urchin sperm flagella and reported to interact with CCDC176 in the mouse sperm axoneme (Leung *et al*., 2023; Liu *et al*., 2023). Together, these observations suggest that association with microtubule-based structures may represent an evolutionarily conserved property of the CCDC81 family. However, whether CCDC81 directly interacts with microtubules, which domains mediate this interaction, and how this activity contributes to centriole organization remains unresolved.

Here, we investigate the molecular properties and cellular function of CCDC81 using *Chlamydomonas reinhardtii* and human cells. By combining U-ExM, biochemical approaches, microtubule dynamics assays, and cryo-electron microscopy, we demonstrate that CCDC81 directly binds and stabilizes microtubules through its N-terminal HU domains. In *Chlamydomonas*, we identify CCDC81 as a component of the A-C linker and characterize its interaction with POC11, the homolog of human CCDC77, revealing a modular organization in which distinct protein domains mediate microtubule binding and protein interactions. Extending these findings to human cells, we show that CCDC81 associates with centrioles and basal bodies and that its depletion compromises centriole architecture and the organization of specific centriolar components, with more pronounced defects upon combined depletion with CCDC77. Together, our findings establish CCDC81 as an evolutionarily conserved microtubule-binding and stabilizing protein, reveal a previously unrecognized function for its HU-related domains, and provide new molecular insights into the mechanisms maintaining centriole structural integrity.

## Results

### CCDC81 and POC11 localize to the A-C linker of *Chlamydomonas reinhardtii* basal bodies

Our previous mass spectrometry analysis of isolated *Chlamydomonas reinhardtii* basal bodies identified CCDC81 as one of the most abundant proteins, ranking first in the proteomic dataset (Hamel *et al*., 2017; POBdb). Interestingly, its human homolog has also been reported to associate with centrioles, including those of sperm cells (Firat-Karalar *et al*., 2014; Liu, Liu and Nai, 2026). These observations prompted us to investigate the precise localization of CCDC81 within *Chlamydomonas* basal bodies. In parallel, we examined POC11, the *Chlamydomonas* homolog of CCDC77, a recently identified component of the A-C linker in human centrioles (Bournonville *et al*., 2025).

To determine the localization of these proteins, we generated polyclonal antibodies against *Chlamydomonas* CCDC81 and POC11. Immunofluorescence analysis of wild-type cells co-stained for tubulin and either CCDC81 or POC11 revealed that both proteins localized to the basal bodies at the base of the flagella (**Figure 1A, B** and **Figure S1A-B**). To validate the specificity of the CCDC81 signal, we analyzed the insertional mutant strain LMJ.RY0402.055207 (**Figure 1A-B, Figure S1C-E**), generated by the *Chlamydomonas* Library Project and hereafter referred to as *ccdc81 m1* (see Materials and Methods). CCDC81 fluorescence at basal bodies was significantly reduced in *ccdc81 m1* compared with wild-type cells, supporting the specificity of the observed signal (**Figure 1A-B**). Interestingly, *ccdc81 m1* basal bodies appeared shorter and displayed structural abnormalities (**Figure S1F, G**). However, because this strain had not been backcrossed and could therefore harbor additional mutations, we could not unambiguously attribute these phenotypes to CCDC81 loss and did not investigate them further. As we did not have a corresponding POC11 mutant strain at the time of this study, we could not perform an equivalent genetic validation of the POC11 antibody. Nevertheless, analysis of POC11 in the *ccdc81 m1* background revealed no detectable change in its basal body signal compared with wild-type cells (**Figure S1A-B**), indicating that POC11 localization to basal bodies is maintained upon CCDC81 depletion.

**Figure 1.**
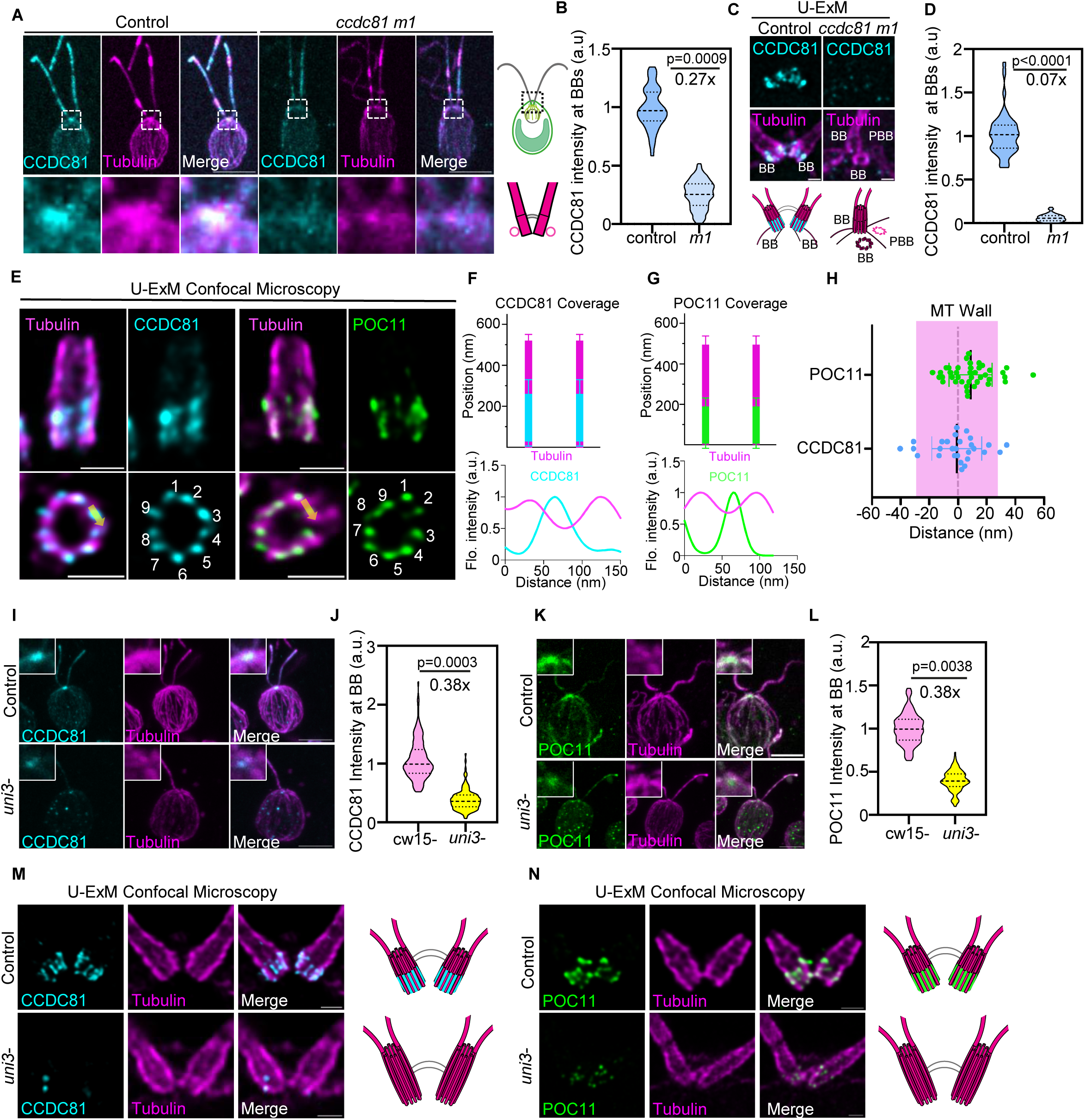
CCDC81 and POC11 localize to the proximal A–C linker region of *Chlamydomonas* basal bodies. N, measurements; n, independent experiments. Signal intensities are normalized to control. Scale bars indicate pre-expansion dimensions. (**A**) Immunofluorescence microscopy of control *Chlamydomonas* cells (left panel) and *ccdc81 m1 mutant* cells (right panel) stained with antibodies against CCDC81 (cyan) and α/β-tubulin (magenta). Dashed boxes indicate the position of the basal bodies, which corresponds to the 4-fold magnified insets below the images. Illustrations of a *Chlamydomonas* cell (top) and basal bodies (bottom) are shown next to the images. Scale bars: 5 μm (images). (**B**) Quantification of CCDC81 normalized signal intensity at basal bodies in control and mutant cells. Control, N = 203, n = 3; *ccdc81 m1*, N = 256, n = 3. Nested t test, two-tailed. (**C**) Expanded *C. reinhardtii* basal bodies stained for α/β-tubulin (magenta) and CCDC81 (cyan), shown in longitudinal view (control) and in longitudinal and top view (*ccdc81 m1*). Basal bodies (BB) and probasal bodies (PBB) are marked, and illustrations of the basal bodies are shown below the images. Scale bar: 1 μm. (**D**) Quantification of CCDC81 normalized signal intensity at basal bodies in control and mutant cells. Control, N = 97, n = 3; *ccdc81 m1*, N = 121, n = 3. Nested t test, two-tailed. (**E**) Isolated *C. reinhardtii* basal bodies expanded and stained for α/β-tubulin (magenta), and CCDC81 (cyan) or POC11 (green). The top panel shows the longitudinal view, and the bottom panel shows the top view. Each triplet microtubule in the bottom panel is from 1 to 9. The yellow arrows placed on the images correspond to the measurement axes of the plot profiles in (F) and (G). Scale bar: 1 μm. (**F, G**) Longitudinal position of CCDC81 (F) and POC11 (G) relative to the tubulin wall (top, n = 3), and the fluorescence intensity of each protein measured along the arrows in (E) relative to the neighbouring microtubule triplets (MTTs) (bottom), demonstrating the precise location of the proteins in between MTTs. (**H**) Radial positions of CCDC81 (N = 28, n = 3) and POC11 (N = 40, n = 3) relative to the tubulin wall, where 0 marks the centre of the tubulin wall. (**I, K**) Immunofluorescence microscopy images of *C. reinhardtii* control cw15-cells (top) or the δ-tubulin mutant *uni3-* (bottom) stained for α/β-tubulin (magenta) and (I) CCDC81 (cyan) or (K) POC11 (green). Insets show the basal body region magnified 4-fold. Scale bar: 5 μm. (**J, L**) The normalized signal intensity of (J) CCDC81 and (L) POC11 at the basal bodies of the *uni3-* strain compared to control. For CCDC81 in cw15-, N = 332, n = 3; uni3-, N = 345, n = 3. For POC11 in cw15-, N = 273, n = 3; uni3-, N = 238, n = 3. Nested t test, two-tailed. (**M, N**) Expanded *C. reinhardtii* cells showing the basal bodies of either control cw15-(top panel) or *uni3-* (bottom) stained for α/β-tubulin (magenta) and (M) CCDC81 (cyan) or (N) POC11 (green). Illustrations summarizing the localization in each strain are shown next to the images. Scale bar: 1 μm.

To determine whether CCDC81 localizes to the A-C linker, we performed U-ExM on *Chlamydomonas* cells (**Figure 1C-D**) and isolated basal bodies (**Figure 1E**). We used POC11, the *Chlamydomonas* homolog of the human A-C linker protein CCDC77, as a reference to define the position of this structure. Both proteins localized to the proximal region of the basal body, extending along 45% and 36.5% of its total length for CCDC81 and POC11, respectively (**Figure 1E–H**). Importantly, top-view analysis revealed that POC11 formed nine discrete foci positioned between adjacent microtubule triplets, consistent with its expected localization at the A-C linker (**Figure 1E–G**). Strikingly, CCDC81 displayed a similar ninefold organization, with signals also positioned between neighboring microtubule triplets (**Figure 1E–G**). Quantitative analysis further showed that both proteins localized at the level of the microtubule wall, with radial positions relative to the tubulin center of mass of −0.80 ± 17.37 nm for CCDC81 and −8.86 ± 14.96 nm for POC11 (**Figure 1H**). Thus, POC11 localization confirms the conserved positioning of this A-C linker component in *Chlamydomonas*, while the similar proximal distribution, ninefold organization and radial positioning of CCDC81 identify it as a previously uncharacterized component of the A-C linker, further supporting the evolutionary conservation of its molecular architecture.

To further investigate whether CCDC81 is associated with the A-C linker, we examined its localization in the *Chlamydomonas uni3-1* mutant, which lacks δ-tubulin and exhibits defects in microtubule triplet assembly, particularly affecting the C-tubule (de Loubresse, Ruiz and Beisson, 2001; Pudlowski *et al*., 2025). Previous structural studies have shown that these defects can lead to partial or complete disruption of the A-C linker (de Loubresse, Ruiz and Beisson, 2001). We therefore reasoned that CCDC81 localization should be affected in this mutant if it is associated with this structure. Using POC11 as a reference A-C linker protein, quantitative immunofluorescence microscopy revealed a marked reduction in the basal body signals of both CCDC81 and POC11 in *uni3-1* compared with control cells (**Figure 1I–L**). To characterize these defects in greater detail, we performed U-ExM (**Figure 1M, N**) and iterative U-ExM (Louvel *et al*., 2023) (iU-ExM; **Figure S2A-C**). Both proteins showed a pronounced disruption of their characteristic proximal localization and ninefold organization, with partial or complete loss of signal in individual basal bodies. Interestingly, these defects were more prominent in mature basal bodies, whereas pro-basal bodies retained CCDC81 and POC11 signals to a greater extent (**Figure S2A-C**). Expansion microscopy also revealed discontinuities in the basal-body microtubule wall (**Figure S2A, D**, white arrows), consistent with the previously reported defects in C-tubule assembly and A-C linker integrity in the *uni3-1* mutant (de Loubresse, Ruiz and Beisson, 2001), together with a modest increase in basal-body length (**Figure S2D, E**). Thus, the disruption of CCDC81 localization in parallel with POC11 following loss of C-tubule integrity provides additional evidence that CCDC81, together with POC11, is associated with the A-C linker and that its proper organization depends on the integrity of the microtubule triplets.

Having established their localization to the A-C linker region, we sought to define more precisely how CCDC81 and POC11 are positioned relative to neighboring microtubule triplets. Taking advantage of the increased spatial resolution provided by iU-ExM, we resolved both proteins between adjacent microtubule triplets in basal bodies and pro-basal bodies *in situ* (**Figure 2A-D; Figure S3A, B**), enabling their relative positions within the A-C linker to be distinguished. Quantitative measurements relative to the A-tubule revealed distinct radial positions for the two proteins, with POC11 positioned closer to the A-tubule and CCDC81 shifted toward the C-tubule (**Figure 2D**). Together, these results establish that CCDC81 and POC11 occupy distinct positions within the A-C linker of *Chlamydomonas* basal bodies.

**Figure 2.**
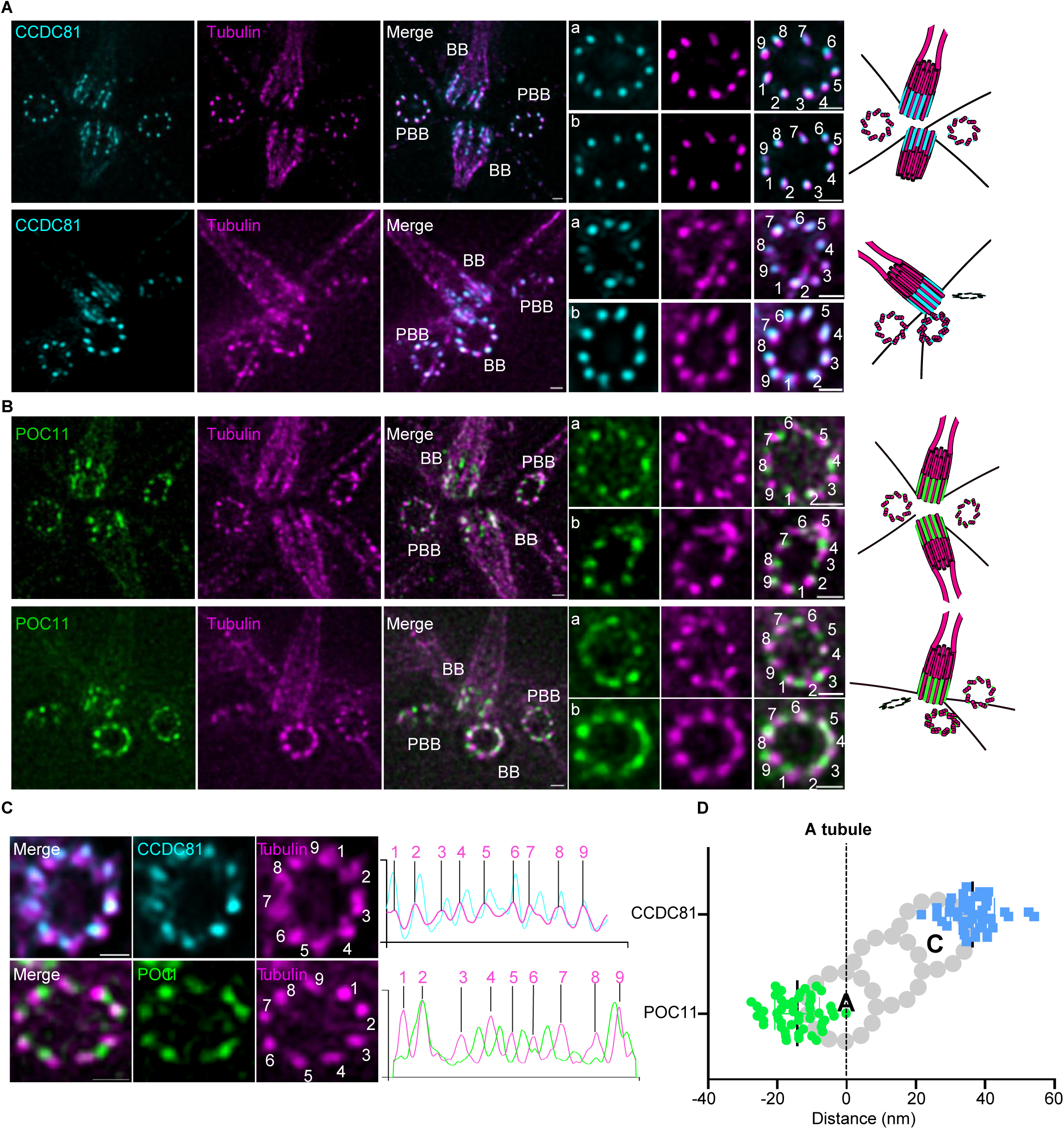
Nanoscale mapping positions CCDC81 and POC11 at distinct sites within the A–C linker. n, independent experiments; N, microtubule triplets measured. Samples were expanded by iterative Ultrastructure Expansion Microscopy (iU-ExM). **(A, B)** Expanded basal bodies of *C. reinhardtii* cells stained for α/β tubulin (magenta), and CCDC81 (cyan, **A**) or POC11 (green, **B**). Basal bodies (BB) and pro-basal bodies (PBB) are marked. Top panels show two PBBs from the top view and two BBs from the lateral view; insets show the PBBs at 4-fold magnification. Bottom panels show one PBB from the top view and one from the side view, while one BB is shown from the side view and the other from the top view; inset (a) shows the PBB and inset (b) the BB, both at 4-fold magnification. MTTs are labelled from 1 to 9. Illustrations summarizing the localization are shown next to the images. Scale bars = 1 μm. **(C)** Expanded basal bodies of *C. reinhardtii* cells shown in top view and stained for α/β tubulin (magenta), and CCDC81 (cyan, top panel) or POC11 (green, bottom panel). Peaks are numbered 1 to 9 to match the MTTs in the corresponding image. Scale bar = 1 μm. **(D)** Distance of CCDC81 and POC11 from the A-tubule. The A-tubule was identified from triplet orientation. Data points are plotted as relative distance from the A-tubule, where minus and plus indicate on which side of the A-tubule the protein sits. A schematic MTT is overlaid for reference, with the A- and C-tubules marked. *n* = 3; CCDC81, *N* = 45; POC11, *N* = 49.

### CCDC81 and POC11 interact with microtubules *in vitro*

Given their localization at the A-C linker and the interface between neighboring microtubule triplets, we next asked whether CCDC81 and POC11 could directly interact with each other and soluble tubulin. To address this, we performed pull-down assays using recombinant MBP-CCDC81 and His-POC11 in the presence or absence of purified tubulin (**Figure 3A-D** and **Figure S4**). His-POC11 efficiently pulled down MBP-CCDC81 and not MBP alone, demonstrating a direct interaction between the two proteins (**Figure 3A**). Moreover, reciprocal pulldown of POC11 with CCDC81 confirmed that this interaction is independent of which protein is used as a bait (**Figure S4**). In addition, these experiments revealed that both CCDC81 and POC11 directly interact with tubulin (**Figure 3B, C** and **Figure S4**). When all three components were combined, CCDC81, POC11, and tubulin were recovered together, indicating that they can assemble into a ternary complex *in vitro* (**Figure 3D** and **Figure S4**).

**Figure 3.**
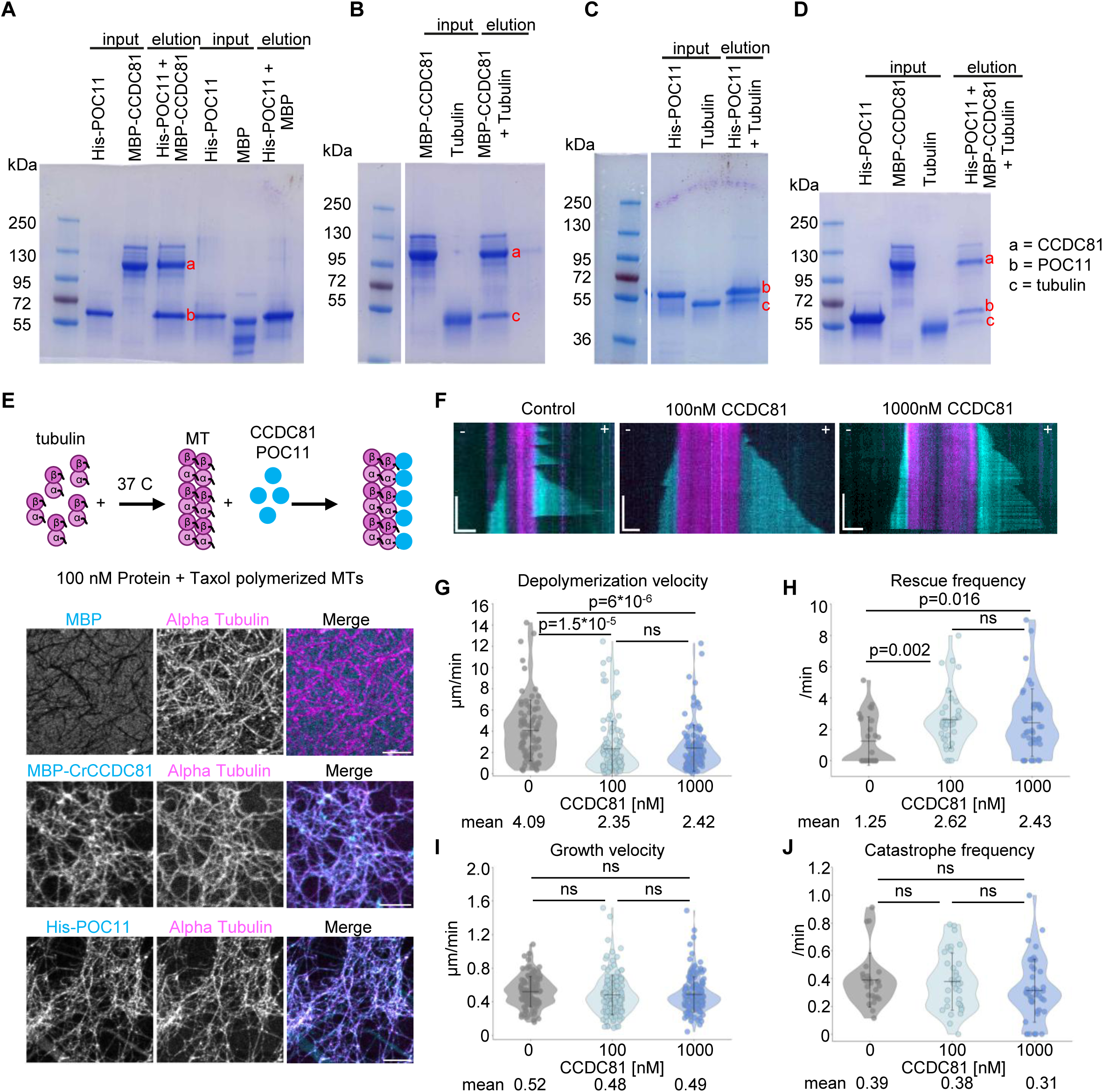
CCDC81 and POC11 interact directly and associate with microtubules *in vitro*. n = 3 independent experiments. For (G–J), *N* is the number of growth or shrinkage events measured for the velocity parameters (G, I), and the number of microtubules analysed for the frequency parameters (H, J). **(A–D)** *In vitro* pulldown assays between: His-POC11 and MBP-CCDC81, or MBP alone as a negative control (A; n = 4 and n = 3 respectively); MBP-CCDC81 and soluble tubulin (B; n = 3); His-POC11 and soluble tubulin (C; n = 3); and His-POC11, MBP-CCDC81 and soluble tubulin (D; n = 4). For the pulldowns in (A), (C) and (D), His-POC11 was immobilized on cobalt resin and then incubated with MBP or MBP-CCDC81 (A), with tubulin (C), or first with MBP-CCDC81 and then with tubulin (D). For the pulldown in (B), MBP-CCDC81 was immobilized on amylose resin and incubated with tubulin. For each gel, the inputs and elutions are shown. Bands are labelled a (MBP-CCDC81), b (His-POC11) and c (tubulin), and molecular weight markers are indicated in kDa. Lanes in (B), (C) and (D) were cropped from the gel, and dividing lines indicate non-adjacent lanes. The CCDC81–POC11 interaction was also confirmed in the reciprocal configuration, with MBP-CCDC81 immobilized on amylose resin instead of His-POC11 on cobalt resin (n = 2; Figure S4), giving six independent observations across two bait configurations. Uncropped gels and replicates are shown in Figure S4**. (E)** *In vitro* microtubule binding assay. The scheme at the top summarizes the experiment. Images are representative of 3 independent experiments. Fixed microtubules were stained for α/β-tubulin (magenta) and MBP, CCDC81 or POC11 (cyan). Scale bar = 5 μm. **(F)** Representative kymographs of microtubule dynamics with 16 µM tubulin in the absence of protein (control) and in the presence of 100 nM or 1000 nM CCDC81. GMPCPP-stabilized microtubule seeds are shown in magenta, and the dynamic GDP microtubule lattice is shown in cyan. Scale bars: horizontal, 1 µm; vertical, 5 min. **(G–J)** Dynamic parameters of GDP microtubule plus ends in the absence of protein (control) and in the presence of 100 nM or 1000 nM CCDC81. **(G)** Depolymerization velocity; control, N = 85; 100 nM, N = 108; 1000 nM, N = 110. **(H)** Rescue frequency; control, N = 27; 100 nM, N = 36; 1000 nM, N = 42. **(I)** Growth velocity; control, N = 106; 100 nM, N = 139; 1000 nM, N = 149. **(J)** Catastrophe frequency; control, N = 25; 100 nM, N = 35; 1000 nM, N = 42. Control is shown in grey, 100 nM CCDC81 in light blue and 1000 nM CCDC81 in blue. The centre line and whiskers represent the mean and S.D. respectively, and the mean value is given below each violin. P values were determined by a two-tailed t test; ns, not significant.

After establishing CCDC81 and POC11 interaction with soluble tubulin, we next asked whether they can interact with polymerized microtubules. To address this, we first performed *in vitro* microtubule co-pelleting assays using recombinant full-length MBP-CCDC81 and His-POC11 purified from *E. coli* (**Figure S5A, B**). Each recombinant protein was incubated either alone or with pre-polymerized microtubules, followed by high-speed centrifugation to separate soluble and microtubule-containing fractions. In the absence of microtubules, a substantial fraction of CCDC81 remained in the soluble fraction, whereas addition of microtubules strongly shifted CCDC81 to the pellet (**Figure S5A**), demonstrating its ability to bind microtubules directly. By contrast, POC11 was recovered almost entirely in the pellet even in the absence of microtubules, indicating limited solubility under these experimental conditions and precluding assessment of its microtubule-binding activity using this assay (**Figure S5B**).

To independently examine the association of CCDC81 and POC11 with microtubules, we next used a microscopy-based approach. Recombinant MBP-CCDC81 and His-POC11 were either incubated with pre-polymerized microtubules (**Figure 3E** and **Figure S5C, D**) or added to soluble tubulin during microtubule polymerization (**Figure S5E, F**), followed by deposition onto coverslips and immunofluorescence imaging. Whereas MBP alone showed no detectable association with microtubules, both CCDC81 and POC11 decorated the microtubule lattice (**Figure 3E; Figure S5C-F**). Notably, CCDC81 copolymerization promoted the formation of thicker microtubule bundles, consistent with its robust association with the microtubule lattice. POC11, in contrast, frequently accumulated in condensate-like structures, which may account for its sedimentation in the absence of microtubules in the co-pelleting assay (**Figure S5D, F**). Consistently, both recombinant proteins decorated microtubules in fixed samples imaged by TIRF microscopy (**Figure S5G, H**). Together, these complementary approaches establish CCDC81 as a direct microtubule-binding protein and support the ability of POC11 to associate with microtubules *in vitro*.

Finally, given the ability of CCDC81 to interact with both soluble tubulin and polymerized microtubules, we next investigated whether it could influence microtubule dynamics. To this end, we immobilized GMPCPP-stabilized microtubule seeds on coverslips and monitored microtubule growth and shrinkage by TIRF microscopy in the presence of 16 µM soluble tubulin and increasing concentrations of CCDC81 (**Figure 3F–J**). Addition of 100 nM CCDC81 significantly reduced the microtubule depolymerization velocity and increased rescue frequency (**Figure 3G, H**). Increasing the CCDC81 concentration to 1 µM did not further enhance these dynamic parameters. By contrast, neither the microtubule growth velocity nor the catastrophe frequency was affected (**Figure 3I, J**). Thus, CCDC81 selectively affects microtubule shrinkage and rescue without altering their growth or catastrophe frequency, indicating that it promotes microtubule stability *in vitro*. Together, these findings show that CCDC81 and POC11 interact directly and can associate with both soluble tubulin and polymerized microtubules. Moreover, they identify CCDC81 as a microtubule-stabilizing protein, providing a potential molecular basis for its association with the A-C linker.

### CCDC81 and POC11 associate with microtubules in cells through distinct domains

Having established that CCDC81 and POC11 associate with microtubules *in vitro*, we next investigated whether they also interact with microtubules in cells. To this end, we expressed GFP-tagged versions of *Chlamydomonas* CCDC81 (CrCCDC81) and POC11 in human U2OS cells and examined their localization along cytoplasmic microtubules (**Figure 4; Figure S6**). GFP-CrCCDC81 robustly decorated the microtubule network in ∼98% of transfected cells and induced prominent microtubule bundling (**Figure 4A**), consistent with the microtubule-binding and bundling activities observed *in vitro*. In contrast, GFP-POC11 decorated cytoplasmic microtubules in fewer than 20% of transfected cells and predominantly accumulated in aggregate-or condensate-like structures that also contained tubulin (**Figure 4B**). We next asked whether association with either protein affected microtubule stability. Following cold-induced microtubule depolymerization, expression of GFP-CrCCDC81 significantly preserved the cytoplasmic microtubule network, whereas GFP-POC11 conferred little or no detectable protection (**Figure S6A-D**). Thus, GFP-CrCCDC81 not only associates with microtubules in cells but also promotes their bundling and protects them against cold-induced depolymerization. This is consistent with its effect *in vitro*, where CCDC81 slowed microtubule depolymerization and increased the rescue frequency (**Figure 3G, H**).

**Figure 4.**
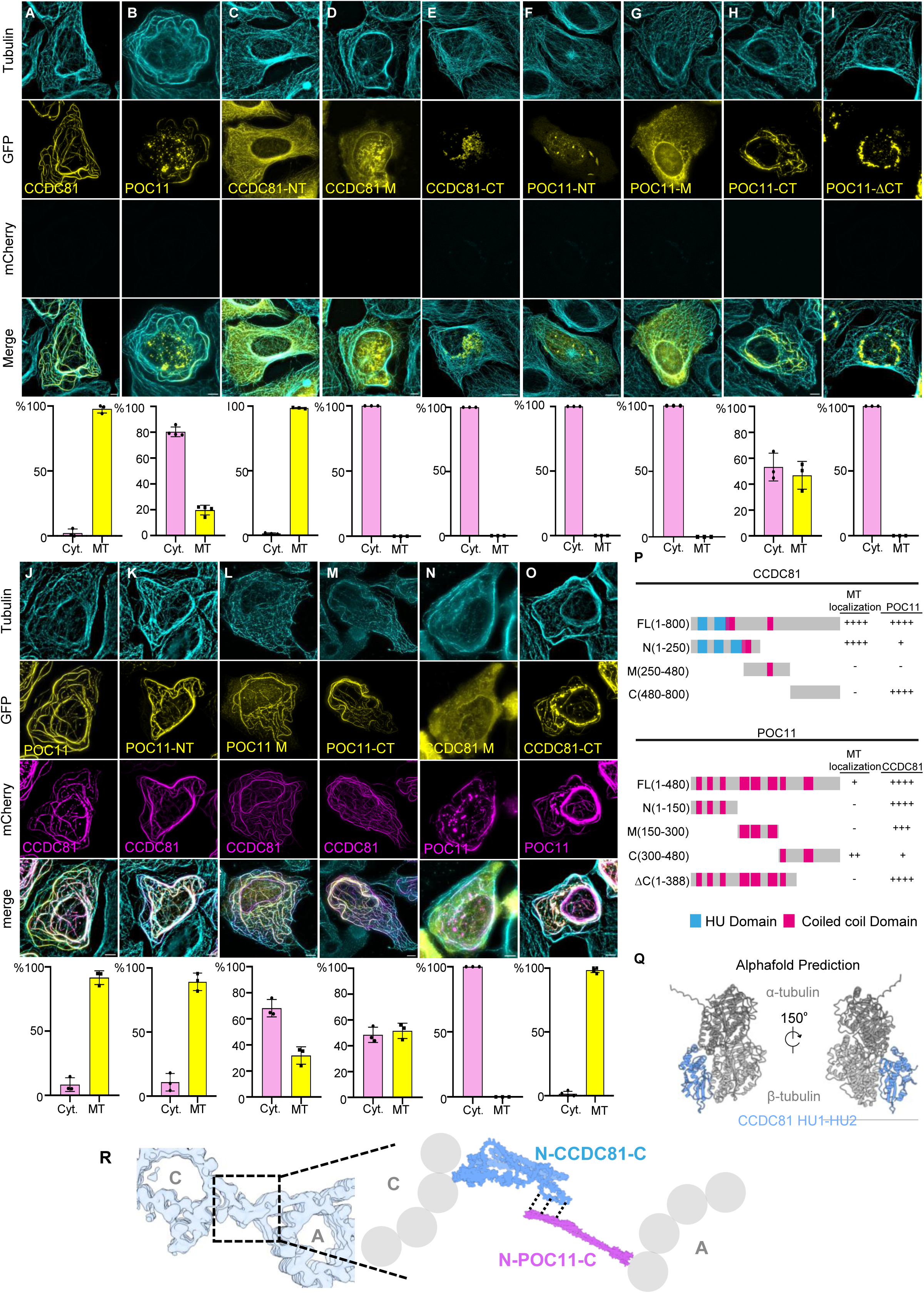
Distinct CCDC81 domains mediate microtubule association and interaction with POC11. **(A–O)** U2OS cells expressing GFP-(yellow) or mCherry-tagged (magenta) *C. reinhardtii* constructs stained for α/β tubulin (cyan). (A), GFP-CrCCDC81 alone. (B) GFP-POC11 alone. (C) GFP-CCDC81 region N terminus. (1-250 aa). (D) GFP-CCDC81 middle region (250-480 aa). (E) GFP-CrCCDC81 C terminus (480-800 aa). (F) GFP-POC11 region N terminus (1-150 aa). (G) GFP-POC11 middle region (150-300 aa). (H) GFP-POC11 C terminus (300-480 aa). (I) GFP-POC11 ΔC-terminus (1-388 aa). (J-M) mCherry-CrCCDC81 co-expressed with GFP-POC11 (J), POC11 N terminus (K), POC11 middle region (L) or POC11 C terminus (M). (N, O) mCherry-POC11 co-expressed with GFP-CrCCDC81 middle region (N) or CCDC81 C terminus (O). *n* is the number of independent experiments and *N* the total number of transfected cells scored: (A) *n* = 3, *N* = 404 (Cyt. 2.14%, MT 97.86%); (B) *n* = 4, *N* = 496 (Cyt. 80.31%, MT 19.69%); (C) *n* = 3, *N* = 349 (Cyt. 1.43%, MT 98.56%); (D) *n* = 3, *N* = 284 (Cyt. 100%, MT 0%); (E) *n* = 3, *N* = 206 (Cyt. 100%, MT 0%); (F) *n* = 3, *N* = 235 (Cyt. 100%, MT 0%); (G) *n* = 3, *N* = 269 (Cyt. 100%, MT 0%); (H) *n* = 3, *N* = 266 (Cyt. 53.20%, MT 46.80%); (I) *n* = 3, *N* = 310 (Cyt. 100%, MT 0%); (J) *n* = 3, *N* = 445 (Cyt. 8.35%, MT 91.65%); (K) *n* = 3, *N* = 284 (Cyt. 10.81%, MT 89.19%); (L) *n* = 3, *N* = 258 (Cyt. 68.16%, MT 31.84%); (M) *n* = 3, *N* = 232 (Cyt. 48.47%, MT 51.53%); (N) *n* = 3, *N* = 79 (Cyt. 100%, MT 0%); (O) *n* = 4, *N* = 124 (Cyt. 1.51%, MT 98.49%). Scale bars = 5 μm. **(P)** Summary of the CCDC81 and POC11 constructs and their localization and interaction properties. HU domains are shown in blue and coiled-coil domains in magenta. Microtubule localization and interaction with the partner protein are scored from − (not detected) to ++++ (strongest interaction). **(Q)** AlphaFold prediction of the CCDC81 N-terminus bound to tubulin. The HU1 and HU2 domains of CCDC81 span residues 1–170. Two views related by a 150° rotation are shown. α- and β-tubulin are colored dark and light grey respectively, and CCDC81 in blue. **(R)** Schematic summarizing the proposed CCDC81–POC11 interaction within the A–C linker. Protein positions are inferred from the localization and interaction data in this figure and are not fitted to density. (Density map from EMD-9174)

We next sought to identify the regions responsible for microtubule association. We generated GFP- or mCherry-tagged constructs encompassing the N-terminal, central, and C-terminal regions of each protein and assessed their association with microtubules in U2OS cells (**Figure 4C-I**, and **Figure 4P**). For CCDC81, microtubule-binding activity mapped to the N-terminal region (amino acids 1–250) (**Figure 4C**), which robustly decorated cytoplasmic microtubules, whereas the central and C-terminal regions showed no detectable association (**Figure 4D, E**). Interestingly, the CCDC81 N-terminus contains two HU (histone-like/heat-unstable) domains followed by a coiled-coil region (CC1) (**Figure 4P** and **Figure S6I**). Further truncation analysis mapped the microtubule-binding activity of CCDC81 to the region encompassing its two HU domains (**Figure S6E-I**). Moreover, AlphaFold modeling predicted a high-confidence interaction between this HU-domain region and β-tubulin (**Figure 4Q; Figure S6N**), providing a potential structural basis for CCDC81-microtubule binding and consistent with the observation of CCDC81-domain proteins within ciliary doublet microtubules (Kubo *et al*., 2023; Legal *et al*., 2023; Gao *et al*., 2024).

Conversely, POC11 microtubule-binding activity mapped to its C-terminal region (amino acids 300–480) (**Figure 4H**), whereas its N-terminal and central regions failed to decorate microtubules (**Figure 4F, G, I**). Further dissection of the C-terminus refined this activity to amino acids 391–481 (**Figure S6J-M**). Thus, CCDC81 and POC11 engage microtubules through distinct regions, with microtubule-binding activity residing within the N-terminal HU domains of CCDC81 and the extreme C-terminus of POC11.

Having identified their respective microtubule-associating regions, we next investigated how CCDC81 and POC11 interact with one another in cells. We took advantage of the robust localization of CrCCDC81 along cytoplasmic microtubules to establish a recruitment assay, in which the redistribution of POC11 to microtubules could serve as a readout of its interaction with CCDC81. Co-expression of GFP-CCDC81 and mCherry-POC11 resulted in their colocalization along cytoplasmic microtubules (**Figure 4J**), consistent with the direct interaction observed *in vitro*. We then used this assay to identify the regions mediating their association. The N-terminal region of POC11, which did not associate with microtubules when expressed alone, was recruited to microtubules in 89% of cells upon co-expression with full-length CCDC81 (**Figure 4K**). In contrast, the central region was recruited in only approximately 30% of cells (**Figure 4L**). The C-terminal region of POC11 was associated with microtubules in approximately 50% of cells when expressed alone, and this proportion remained unchanged upon co-expression with CCDC81 (**Figure 4M**). These observations indicate that the N-terminal region of POC11 mediates its interaction with CCDC81, whereas its C-terminal region can associate with microtubules independently.

We next performed the reciprocal recruitment assay to identify the region of CCDC81 involved in POC11 binding. The C-terminal region of CCDC81, which did not associate with microtubules when expressed alone, was recruited to microtubules upon co-expression with POC11, whereas its central region was not (**Figure 4N, O**). This identifies the C-terminus of CCDC81 as the principal region mediating its association with POC11. Guided by these experimental observations, we used AlphaFold to model the interaction between the C-terminal region of CCDC81 (amino acids 747–800) and the N-terminal region of POC11 (amino acids 1–51). The resulting model predicted an interaction between these regions (**Figure S6O**), consistent with the recruitment assays and suggesting a possible molecular interface for complex formation. Together, these results reveal a modular organization of the CCDC81–POC11 complex, in which the N-terminal region of CCDC81 and the C-terminal region of POC11 mediate microtubule association, while the C-terminal region of CCDC81 and the N-terminal region of POC11 mediate their interaction (**Figure 4R**).

### CCDC81 associates with the outer surface of the microtubule lattice

To determine how CCDC81 associates with microtubules, we investigated whether it binds to the outer microtubule surface or behaves as a microtubule inner protein (MIP) (Kubo *et al*., 2023; Legal *et al*., 2023; Gao *et al*., 2024), as previously described for other microtubule-associated proteins (Ichikawa and Bui, 2018; Owa *et al*., 2019). To distinguish between these possibilities, we incubated taxol-stabilized microtubules with purified MBP-CrCCDC81 and analyzed them by cryo-electron microscopy (cryo-EM) (Figure 5A). Compared with undecorated microtubules, those incubated with MBP-CrCCDC81 exhibited additional, irregular densities along their outer surface (Figure 5B, C). Power spectrum analysis revealed a prominent layer line at approximately 4 nm, corresponding to the axial repeat of tubulin, but no additional periodic features attributable to CCDC81. This suggests that CCDC81 either associates with the microtubule lattice without a well-defined periodic arrangement or follows the underlying tubulin periodicity.

**Figure 5.**
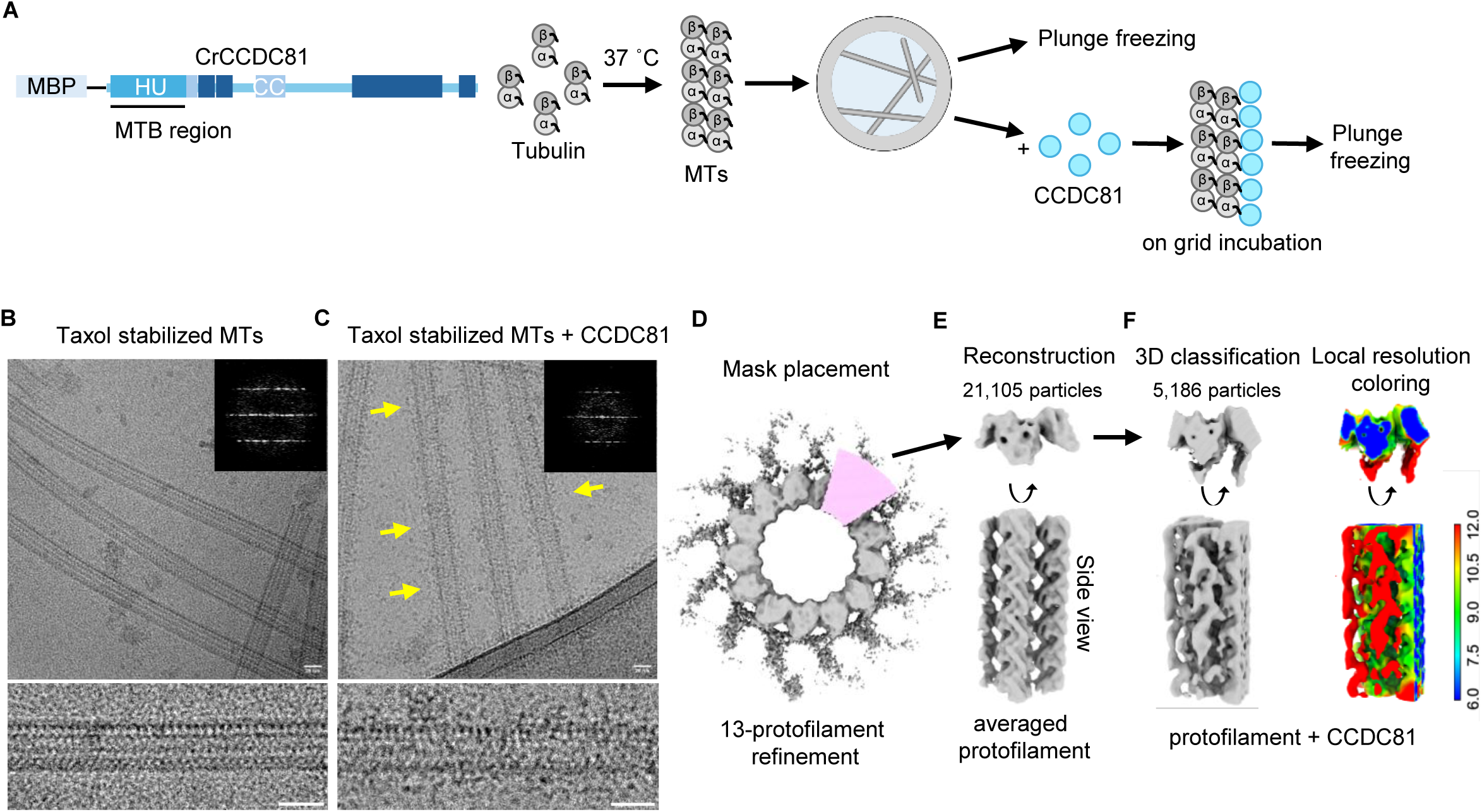
CCDC81 associates with the outer microtubule surface. **(A)** Experimental design. Left, domain organization of MBP-CrCCDC81 is shown and microtubule binding region is indicated. Right, tubulin was polymerized into Taxol-stabilized microtubules and applied to cryo-EM grids. Grids were either plunge-frozen directly (undecorated microtubules) or incubated with MBP-CrCCDC81 on the grid before plunge-freezing. **(B)** Taxol-stabilized microtubules alone. Top, cryo-EM micrograph; inset, power spectrum of a straightened microtubule, showing the layer lines of the tubulin lattice. Bottom, a straightened microtubule. **(C)** As in B, for Taxol-stabilized microtubules incubated with MBP-CrCCDC81. Yellow arrows show the irregular extra density visible along the microtubule walls. **(D)** Single-particle processing of MBP-CrCCDC81 incubated microtubules (workflow in Figure S7). Top view of 13-protofilament helical refinement, with the mask (pink) used to isolate one protofilament for signal subtraction and local refinement. (E) Local refinement of one protofilament after signal subtraction (21,105 particles from the merged 12- and 13-protofilament branches; 6.7 Å, Figure S7). Top and side views. (F) 3D classification of the particles in E into four classes. Class 0 (5,186 particles; 7.6 Å, Figure S7G) is shown in top and side views (left) and colored by local resolution (right; key in Å). Scale bars: 25 nm.

To further characterize this interaction and determine whether the additional density could be resolved by averaging, we processed the cryo-EM images using the MiCSPARC workflow implemented in cryoSPARC (Punjani *et al*., 2017; Zhang *et al*., 2026) (Figure S7A, B). Most microtubule segments contained either 12 or 13 protofilaments (Figure S7C), and both populations were independently refined (Figure S7D, E). Following symmetry expansion and signal subtraction to isolate individual protofilaments (Figure 5D, E), local refinement of 21,105 particles yielded a reconstruction of the tubulin lattice at 6.7 Å resolution (Figure 5E and S7F). Although the tubulin lattice was well resolved, no additional density corresponding to CCDC81 could be unambiguously identified in the averaged reconstruction. We therefore performed 3D classification, which revealed an additional density extending along the outer surface of the protofilament, potentially corresponding to bound CCDC81 (Figure 5F and Figure S7G). This observation is consistent with the external decoration observed in the raw cryo-EM images (Figure 5C). However, the limited definition of this density, potentially reflecting conformational flexibility or heterogeneous binding of CCDC81, prevented us from determining its precise binding site, orientation, or stoichiometry on the microtubule lattice. Together, these observations support an association of CCDC81 with the outer microtubule surface rather than its incorporation as a MIP, although its molecular arrangement remains unresolved.

### Human CCDC81 localizes to centrioles and contributes to centriole architecture

Having identified CCDC81 and POC11 as components of the A-C linker in *Chlamydomonas*, we next investigated whether their localization and function are conserved in vertebrates. CCDC77, the human homolog of POC11, was recently identified as a component of the human A-C linker (Bournonville *et al*., 2025), whereas the precise localization and function of human CCDC81 remain less well characterized. We compared CCDC81 sequences from *C. reinhardtii*, *S. purpuratus*, *X. laevis*, *M. musculus*, and *H. sapiens* (**Figures S8, S9**). Human CCDC81 (HsCCDC81) is shorter than its *Chlamydomonas* counterpart (652 versus 800 residues) but shares a similar domain organization, comprising two N-terminal HU domains followed by disordered and coiled-coil regions (**Figure 6A**). Sequence conservation was highest within the HU domains (**Figure 6B**; **Figures S8, S10B**), which were predicted with high confidence by AlphaFold and showed closely superimposable structures between the two species (**Figure S10A, C**). A similar fold was predicted for the N-terminal region of *Tetrahymena* CCDC81 (UniProt ID: I7MLF6), previously proposed to mediate microtubule binding (Legal *et al*., 2023) (**Figure S10D**). Thus, despite sequence divergence, the overall structural organization of CCDC81 appears conserved across evolution.

**Figure 6.**
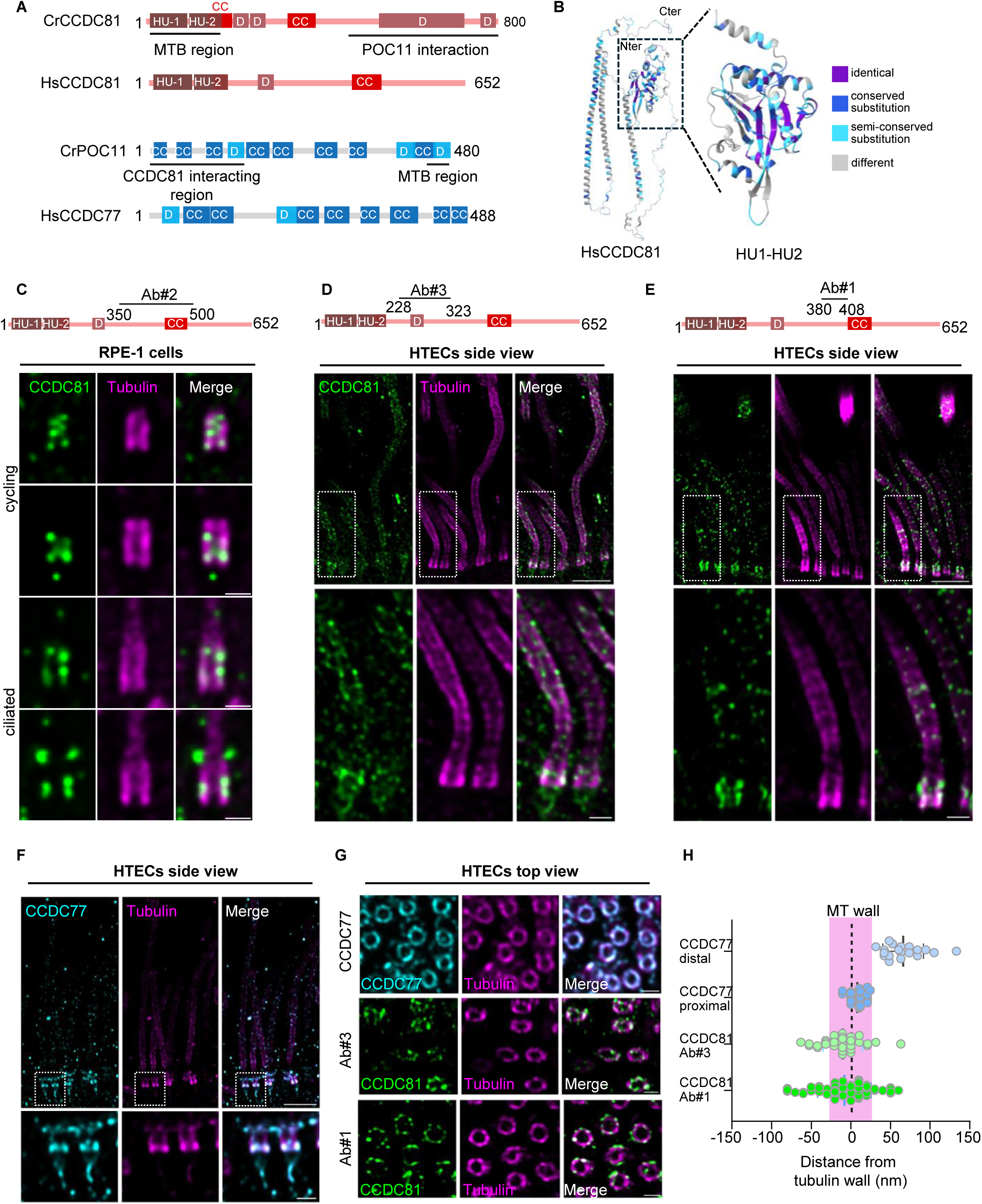
HsCCDC81 and CCDC77 localize to centrioles and basal bodies in human cells. **(A)** Domain organization of *C. reinhardtii* CCDC81 (CrCCDC81, 800 residues) and its human orthologue HsCCDC81 (652 residues), and of *C. reinhardtii* POC11 (CrPOC11, 480 residues) and its human orthologue HsCCDC77 (488 residues). HU-1 and HU-2, HU-like domains; CC, predicted coiled-coil; D, predicted disordered segment. Numbers are residues. The microtubule-binding regions and the CCDC81– POC11 interacting regions mapped in Figures 4 and S6 are indicated beneath the *Chlamydomonas* proteins. **(B)** AlphaFold model of HsCCDC81 colored by sequence conservation, with the N and C termini indicated. The boxed region containing HU-1 and HU-2 is enlarged on the right. Conservation was scored in Jalview (version 2.7) from a multiple sequence alignment of CCDC81 orthologues from five species; the alignment and the per-residue conservation scores are shown in Figure S8. **(C)** Cycling and ciliated RPE-1 cells processed for U-ExM and stained for CCDC81 (green, Ab#2) and α-tubulin (magenta). The region recognized by Ab#2 (residues 350–500) is mapped on the HsCCDC81 diagram above. Four independent CCDC81 antibodies were tested; all label centrioles and basal bodies, and the remaining three are shown in Figure S11. Scale bars = 1 µm. **(D)** Human tracheal epithelial cells (HTECs) differentiated at the air–liquid interface, processed for U-ExM and stained for CCDC81 (green, Ab#3; residues 228–323) and α-tubulin (magenta). 4X enlarged insets shown below. Scale bars = 5 µm (upper) and 1 µm (lower). **(E)** HTECs prepared as in (D) and stained with a second, independent CCDC81 antibody (green, Ab#1; residues 380–408). Scale bars = 5 µm (upper) and 1 µm (lower). **(F)** HTECs prepared as in (D) and stained for CCDC77 (cyan) and α-tubulin (magenta). CCDC77 occupies two positions along the centriole: one in the proximal region, where the A–C linker is located, and one in the distal region, at the level of the appendages. Scale bars = 5 µm (upper) and 1 µm (lower). **(G)** Top views of HTEC centrioles stained for α-tubulin (magenta) together with CCDC77 (cyan) or with CCDC81 (green) detected using Ab#3 or Ab#1. Scale bars = 1 µm. **(H)** Distance of the CCDC77 and CCDC81 signals from the tubulin wall, measured on top views as in (G). Each point is one centriole. The dashed line marks the center of the tubulin wall. Positive values lie outside the wall and negative values towards the lumen. CCDC77 was scored separately at its proximal and distal positions. *N* = 19 centrioles for CCDC77 distal, *N* = 35 for CCDC77 proximal, *N* = 36 for CCDC81 Ab#3 and *N* = 64 for CCDC81 Ab#1. All distances are corrected for the expansion factor (4X).

We next examined the localization of endogenous HsCCDC81 in human RPE-1 cells, using four independent antibodies targeting distinct regions of the protein (**Figure 6C-E**; **Figure S11**). By immunofluorescence microscopy, three out of the four antibodies (Abs1, 2, 3) labelled the centrosome, where the signal overlapped with centriolar marker centrin-2, albeit with some variabilities in the background level and across the cell cycle notably during mitosis (**Figure S11A-D**). This agrees with previous observations using tagged HsCCDC81 (Firat-Karalar *et al*., 2014; Liu, Liu and Nai, 2026). By U-ExM, all four antibodies labelled centrioles (**Figure 6C**; **Figure S11E-G**), in both cycling and ciliated cells (**Figure 6C**). However, unlike CrCCDC81, which localizes to the A-C linker, HsCCDC81 displayed a discontinuous, punctate distribution along the centriolar microtubule wall, without a clear restriction to a specific substructure (**Figure 6C; Figure S11E–G**). Thus, although centriolar localization of CCDC81 is conserved, its nanoscale organization appears to differ between *Chlamydomonas* and human cells.

We further investigated HsCCDC81 in human tracheal epithelial cells (HTECs), in which numerous basal bodies template motile cilia, and compared it with CCDC77. Using two independent antibodies, HsCCDC81 was detected at basal bodies with a punctate distribution and, in addition, along motile cilia (**Figure 6D, E**), consistent with the ciliary localization of CCDC81-family proteins reported in other model systems (Kubo *et al*., 2023; Leung *et al*., 2023; Liu *et al*., 2023; Gao *et al*., 2024, 2024). In contrast, CCDC77 localized to two distinct regions of the basal body: a proximal pool corresponding to the A-C linker region and a distal pool near the appendages (**Figure 6F**), consistent with previous observation in RPE1 cells (Bournonville *et al*., 2025). To refine the radial localization of both proteins, we measured the distance of each signal from the center of the tubulin wall in top views (**Figure 6G, H**). The proximal pool of CCDC77 was located at the level of the wall (7.96 ± 8.98 nm), whereas the distal pool was located outside (65.97 ± 25.50 nm). With both antibodies the HsCCDC81 signal was centered on the microtubule wall (-8.14 ± 33.76 nm for Ab#1 and -10.22 ± 24.55 nm for Ab#3). These findings further support their conserved association with the centriolar structures in ciliated cells.

Having established the conserved association of CCDC81 with centrioles and basal bodies, we next investigated its contribution to centriole function and architecture. Previous work reported reduced ciliogenesis and shorter primary cilia following CCDC81 depletion (Liu, Liu and Nai, 2026). Under our experimental conditions, HsCCDC81 depletion did not significantly affect the proportion of ciliated cells, although primary cilia were longer than in control cells (**Figure S12A-C**). However, U-ExM revealed pronounced structural abnormalities of centrioles reaching 54% following HsCCDC81 depletion compared with 9% in control cells (**Figure 7A, B; Figure S12D, E**). These observations prompted us to investigate in greater detail how loss of HsCCDC81 affects centriole architecture.

**Figure 7.**
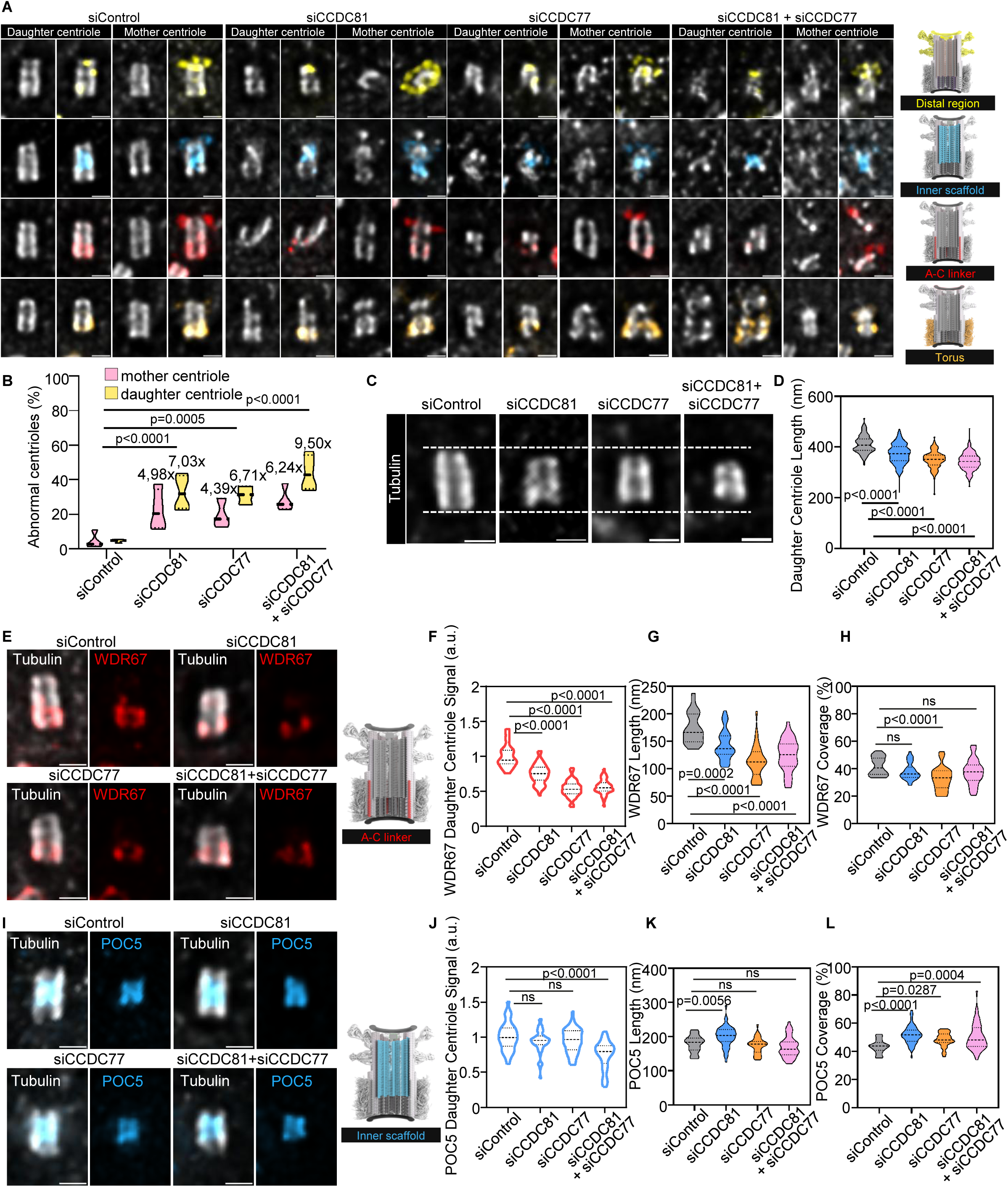
CCDC81 depletion compromises centriole architecture and A–C linker organization. **(A)** siRNA transfected RPE-1 cells processed for U-ExM and stained for α/β-tubulin (grey) together with a marker for each centriolar region: C2CD3 for the distal region (yellow), POC5 for the inner scaffold (cyan), WDR67 for the A–C linker (red) and CEP63 for the torus (orange). CEP164 was co-stained in all experiments to identify the mother centriole. For each condition, the daughter and the mother centriole are shown as tubulin alone and as the merge. The schematics on the right indicate the position of each region within the centriole. Scale bars = 1 µm. **(B)** Percentage of abnormal mother (pink) and daughter (yellow) centrioles in each condition. Centrioles were scored as abnormal when the tubulin signal departed from the regular cylinder seen in controls: gaps in the wall, missing tubulin, twisting, bending, splaying or reduced diameter. The values above each violin are the fold change relative to the corresponding siControl centriole. For siControl: *N* = 473, *n* = 4 for mother centriole (mean = 4.49%), *N* = 476, *n* = 4 for daughter centriole (mean = 4.62%). For siCCDC81: *N* = 704, *n* = 4 for mother centriole (mean = 22.38%), *N* = 723, *n* = 4 for daughter centriole (mean = 32.51%). For siCCDC77: *N* = 374, *n* = 3 for mother centriole (mean = 19.71%), *N* = 398, *n* = 3 for daughter centriole (mean = 31.03%). For siCCDC81 + siCCDC77: *N* = 597, *n* = 4 for mother centriole (mean = 28.04%), *N* = 604, *n* = 4 for daughter centriole (mean = 43.91%). P values were determined by two-way ANOVA with Tukey’s post-hoc test. **(C)** Representative daughter centrioles stained for α/β-tubulin. The dashed lines are drawn at the same position in all four images and mark the extent of the control daughter centriole. Scale bars = 1 µm. **(D)** Daughter centriole length measured from the tubulin signal under the indicated siRNA conditions. For siControl: *N* = 265, *n* = 3 (mean = 410.9 nm). For siCCDC81: *N* = 506, *n* = 3 (mean = 372.5 nm). For siCCDC77: *N* = 337, *n* = 3 (mean = 350.5 nm). For siCCDC81 + siCCDC77: *N* = 455, *n* = 3 (mean = 343.0 nm). P values were determined by one-way ANOVA with Dunnett’s post-hoc test. **(E)** Daughter centrioles stained for α/β-tubulin (grey) and WDR67 (red); for each condition, the merge and the WDR67 channel alone are shown. The schematic indicates the position of the A–C linker. Scale bars = 1 µm. **(F)** WDR67 signal intensity on the daughter centriole, normalized to siControl. For siControl: *N* = 65, *n* = 3 (mean = 1). For siCCDC81: *N* = 140, *n* = 3 (mean = 0.76). For siCCDC77: *N* = 66, *n* = 3 (mean = 0.58). For siCCDC81 + siCCDC77: *N* = 85, *n* = 3 (mean = 0.63). **(G)** The length of the WDR67 signal along the centriole. For siControl: *N* = 60, *n* = 3 (mean = 174.7 nm). For siCCDC81: *N* = 89, *n* = 3 (mean = 149.5 nm). For siCCDC77: *N* = 61, *n* = 3 (mean = 135.6 nm). For siCCDC81 + siCCDC77: *N* = 86, *n* = 3 (mean = 140.7 nm). **(H)** WDR67 coverage, the fraction of the centriole length occupied by the WDR67 signal. For siControl; *N* = 60, *n* = 3 (mean = 40.92%). For siCCDC81; *N* = 89, *n* = 3 (mean = 38.53%). For siCCDC77: *N* = 61, *n* = 3 (mean = 36.79%). For siCCDC81 + siCCDC77: *N* = 86, *n* = 3 (mean = 39.32%). **(I)** Daughter centrioles stained for α/β-tubulin (grey) and POC5 (cyan), shown as in (E). The schematic indicates the position of the inner scaffold. Scale bars = 1 µm. **(J–L)** POC5 signal intensity (J), length (K) and coverage (L), measured and displayed as in (F–H). (J) For siControl: *N* = 64, *n* = 3 (mean = 1). For siCCDC81: *N* = 135, *n* = 3 (mean = 0.94). For siCCDC77: *N* = 98, *n* = 3 (mean = 0.95). For siCCDC81 + siCCDC77: *N* = 143, *n* = 3 (mean = 0.74). (K) For siControl: *N* = 58, *n* = 3 (mean = 178.9 nm). For siCCDC81: *N* = 128, *n* = 3 (mean = 200.1 nm). For siCCDC77: *N* = 93, *n* = 3 (mean = 173.4 nm). For siCCDC81 + siCCDC77: *N* = 138, *n* = 3 (mean = 169.7 nm). (L) For siControl: *N* = 58, *n* = 3 (mean = 43.50%). For siCCDC81: *N* = 128, *n* = 3 (mean = 51.63%). For siCCDC77: *N* = 93, *n* = 3 (mean = 48.26%). For siCCDC81 + siCCDC77: *N* = 138, *n* = 3 (mean = 50.19%). Violin plots show the full distribution with the median and quartiles indicated. *n* = number independent experiments and *N* = number centrioles per condition. P values in **(F-H, J-L)** were determined by one-way ANOVA with Dunnett’s post-hoc test. Measurements are corrected for expansion factor (4X). Knockdown efficiency and the remaining markers and quantifications are shown in Figure S12.

We therefore examined mother (MC) and daughter centrioles (DC) following individual or combined depletion of HsCCDC81 and CCDC77, given that newly assembled daughter centrioles are often more susceptible to siRNA-mediated perturbations (Bournonville *et al*., 2025). Quantitative analysis confirmed efficient depletion of both proteins under our experimental conditions (**Figure S12F**, **G** and **J**, **K**). Whereas control centrioles displayed continuous, elongated microtubule walls, HsCCDC81 depletion led to an increase of abnormal centrioles, including shortened, distorted, and apparently discontinuous structures (MC: 22%; DC: 32.5% compared to 4.5% in control) (**Figure 7A, B; Figure S12D, E**). A similar defect was previously observed following CCDC77 depletion (MC: 18%; DC: 31%) (Bournonville *et al*., 2025) and was further enhanced upon co-depletion of HsCCDC81 and CCDC77, particularly in daughter centrioles (MC: 28%; DC: 44%) (**Figure 7A, B; Figure S12D, E**). Consistent with these morphological abnormalities, measurement of the tubulin signal revealed a slight reduction in daughter centriole length following depletion of either protein, with the strongest effect observed upon their combined depletion (**Figure 7C, D**). These observations indicate that HsCCDC81 and CCDC77 contribute to centriole architecture.

We next investigated whether these morphological defects were accompanied by alterations in specific centriolar structural modules, focusing first on the A-C linker. We examined the interdependence of HsCCDC81 and CCDC77, a conserved A-C linker component and found that CCDC77 depletion impaired the centriolar localization of HsCCDC81, whereas HsCCDC81 depletion did not noticeably affect CCDC77 localization (**Figure S12F, G, J–M**). This asymmetric dependency suggests that CCDC77 may contribute to the recruitment or maintenance of HsCCDC81 at centrioles. To determine whether HsCCDC81 depletion also affects other A-C linker components, we examined WDR67 (Bournonville *et al*., 2025). Interestingly, loss of HsCCDC81 reduced both the fluorescence intensity and longitudinal extent of WDR67 at daughter centrioles (**Figure 7E-H**). These defects were more pronounced following CCDC77 depletion, while combined depletion similarly compromised WDR67 organization (**Figure 7E-H**). Together, these findings reveal a functional interplay between CCDC81 and A-C linker components in human centrioles, despite HsCCDC81 itself not being specifically restricted to the A-C linker region.

We next examined POC5, a component of the centriolar inner scaffold, to assess whether the defects extended beyond the A-C linker. POC5 levels were not affected following individual depletion of HsCCDC81 or CCDC77 but decreased upon co-depletion (**Figure 7I–L**). In contrast, its longitudinal distribution was relatively preserved, suggesting that the inner scaffold is less sensitive than the A-C linker to the loss of either protein. We further examined C2CD3, a distal centriolar component, and CEP63, a component of the proximal torus (**Figure S12H, I**). C2CD3 levels were moderately reduced under all depletion conditions (**Figure S12H, I**), whereas CEP63 showed reduced intensity and longitudinal extent, particularly following CCDC77 depletion, as previously reported (Bournonville *et al*., 2025), and to a lesser extent following HsCCDC81 depletion (**Figure S12N–Q**).

Together, these results establish HsCCDC81 as a contributor to human centriole architecture. Although its localization doesn’t appear completely restricted to the A-C linker, HsCCDC81 depletion strongly affects the organization of this structure, while also perturbing other centriolar components. The enhanced defects observed upon combined depletion of HsCCDC81 and CCDC77 further suggest that these proteins cooperate to maintain centriole structural integrity, supporting a conserved functional relationship between CCDC81 and the A-C linker.

## Discussion

The remarkable stability and ninefold organization of centrioles rely on an intricate protein network associated with the microtubule wall, yet the molecular mechanisms underlying their structural integrity remain incompletely understood. CCDC81 was initially identified through proteomic analysis of mammalian sperm centrioles (Firat-Karalar *et al*., 2014) and subsequently detected as a prominent component of the *Chlamydomonas* basal body proteome (Hamel *et al*., 2017). Despite these observations, its precise localization and molecular function at centrioles remained largely unexplored. Here, we establish CCDC81 as a microtubule-binding protein contributing to centriole architecture and identify POC11 as one of its molecular partners. Their association with the proximal A-C linker in *Chlamydomonas*, together with the structural defects observed following CCDC81 depletion in human cells, suggests that CCDC81 contributes to centriole architecture across evolution, despite some differences in its precise localization. More broadly, our findings provide insights into the function of CCDC81 in participating in the organization and maintenance of the centriolar architecture.

The centriole wall is organized by multiple molecular assemblies, including the cartwheel, A-C linker, inner scaffold, distal luminal ring, or the distal cap, all contributing to centriole integrity (Dobbelaere *et al*., 2020; Le Guennec *et al*., 2020; Steib *et al*., 2020; Bertiaux *et al*., 2025; Bournonville *et al*., 2025). Our findings suggest that CCDC81 contributes to this overall organization through both direct microtubule binding and interactions with other structural proteins. In *Chlamydomonas*, CCDC81 associates with POC11, the homolog of human CCDC77, and their complementary microtubule-binding and interaction domains suggest a possible mechanism for connecting neighboring microtubule triplets. This organization is consistent with their localization at the A-C linker, although the precise arrangement of the complex within the native structure remains to be determined. Importantly, CCDC81 also promotes microtubule bundling and resistance to cold-induced depolymerization, suggesting that its structural contribution may extend beyond simply recruiting other proteins to the centriole wall. These properties suggest that CCDC81 may contribute to centriole integrity through both direct interactions with the microtubule lattice and its association with other structural components. Whether its microtubule-stabilizing activity observed in experimental assays also contributes to centriole stability *in vivo* remains to be established. In human cells, CCDC81 depletion leads to pronounced centriole architectural defects, particularly affecting daughter centrioles. These defects are accompanied by alterations in WDR67 and other centriolar markers, whereas the inner scaffold component POC5 is comparatively less affected. This differential response suggests that CCDC81 contributes to the organization of specific structural modules rather than uniformly affecting the entire centriole architecture. Moreover, combined depletion of CCDC81 and CCDC77 exacerbates centriole abnormalities, supporting complementary contributions of these proteins to centriole integrity. The pronounced defects observed in daughter centrioles may reflect their greater depletion of CCDC81 under our experimental conditions, although a particular requirement for CCDC81 during centriole assembly or maturation cannot be excluded. Importantly, the broader distribution of CCDC81 along human centrioles, compared with its more restricted localization in *Chlamydomonas*, suggests that its structural contribution may not be limited to the A-C linker. Whether CCDC81 associates with distinct molecular complexes along the human centriole wall, and how these interactions contribute to centriole integrity, remain important questions.

An intriguing feature of CCDC81 is the presence of two N-terminal HU-like domains, evolutionarily related to bacterial HU proteins involved in DNA organization. Previous evolutionary analyses proposed that these divergent domains may have acquired alternative interaction partners during eukaryotic evolution (Burroughs *et al*., 2017). Although CCDC81-domain proteins had previously been observed in association with ciliary doublet microtubules (Kubo *et al*., 2023; Legal *et al*., 2023; Gao *et al*., 2024), our data provide direct evidence that the HU-containing region is sufficient for microtubule binding, supporting this hypothesis and suggesting that an ancestral nucleic acid-associated fold may have been repurposed for cytoskeletal organization. This raises the question of how HU domains recognize the tubulin lattice and whether their interaction involves structural features inherited from ancestral HU proteins or newly evolved binding interfaces. Our AlphaFold predictions suggest a potential interaction between the HU-containing region of CCDC81 and β-tubulin, although the molecular determinants of this association remain to be experimentally established. Whether the two HU domains cooperate in microtubule recognition or have acquired distinct functions also remains unknown. More broadly, the ability of CCDC81 to bind and bundle microtubules illustrates how ancient protein folds may have diversified to perform specialized functions within the eukaryotic cytoskeleton.

Interestingly, CCDC81-related proteins appear to associate with distinct microtubule architectures. CCDC81B has been proposed as a microtubule inner protein in *Tetrahymena* ciliary doublets (Gao *et al*., 2024), whereas our cryo-EM observations support an association of *Chlamydomonas* CCDC81 with the outer microtubule surface. However, because the protein was added to pre-assembled microtubules, our experiments do not completely exclude binding inside the lumen. Moreover, while *Chlamydomonas* CCDC81 is concentrated in the proximal basal body, human CCDC81 displays a broader distribution along centrioles and is also detected in motile cilia. These findings suggest that microtubule association may represent a conserved property of the CCDC81 family, whereas its precise localization and molecular partners have diversified across eukaryotes. Thus, CCDC81 may contribute to different microtubule-associated assemblies, with its localization at the A-C linker representing one of several structural contexts. Whether these distinct distributions reflect evolutionary specialization or different interactions of CCDC81 family proteins with the microtubule lattice remains to be determined.

The contribution of CCDC81 to centriole architecture also raises the question of how structural integrity influences ciliogenesis. Previous work reported reduced ciliogenesis and shorter primary cilia following CCDC81 depletion (Liu, Liu and Nai, 2026). In contrast, under our experimental conditions, CCDC81 depletion did not substantially affect ciliogenesis frequency but resulted in longer cilia. Although these differences may reflect variations in depletion efficiency, experimental timing, or cellular context, they may also relate to the differential sensitivity of mother and daughter centrioles to CCDC81 depletion. Indeed, we observed that structural defects predominantly affected daughter centrioles, whereas mother centrioles were comparatively preserved. Since primary cilia assemble from mature mother centrioles, this could explain why the pronounced centriolar abnormalities observed upon CCDC81 depletion were not accompanied by a reduction in ciliogenesis. Thus, the impact of CCDC81 depletion on centriole architecture may not directly translate into defects in cilium formation, particularly when the mother centriole remains sufficiently intact to function as a basal body. Nevertheless, the increased cilium length suggests that CCDC81 may influence additional aspects of ciliary biology. Whether this phenotype reflects subtle alterations in basal body organization or a more direct role of CCDC81 in ciliary microtubule regulation, consistent with its localization along motile cilia, remains to be determined.

Together, our findings establish CCDC81 as a microtubule-associated protein contributing to centriole architecture and reveal a molecular connection between microtubule binding and the organization of centriolar structural assemblies. The evolutionary conservation of its HU domains, together with its distinct localization across centrioles and cilia, suggests that CCDC81-related proteins have diversified to perform structural functions in different microtubule-based organelles. These findings provide a foundation for understanding how CCDC81 contributes to the assembly, stability and functional organization of centrioles.

## Methods

### Chlamydomonas strains

The *Chlamydomonas* strains used in this study were grown in liquid Tris-acetate-phosphate medium (TAP medium containing Trace) at **∼**22°C or on TAP plates with 1.5% agar (Gorman and Levine, 1965). The *Chlamydomonas* mutant, LMJ.RY0402.055207 (*ccdc81* m1) was generated by the CLIP library group (Li *et al*., 2016; X. Li *et al*., 2019) and obtained through the *Chlamydomonas* Resource Center (http://www.chlamycollection.org/). Upon receipt, the mutant was streaked to single colonies, maintained at room temperature on TAP plates, and the insertion site of the CIB1 cassette verified by PCR. The *uni-3-CC4179* mutant strain was obtained from the *Chlamydomonas* Resource Center.

### Isolation of *Chlamydomonas* centrioles

*Chlamydomonas* centrioles were isolated as previously described (Klena *et al*., 2018). Briefly, *cw15-Chlamydomonas reinhardtii* cells were grown to ∼1 x 107 cells/mL density in TAP media. Cells were then deflagellated with pH shock in 10 mM HEPES containing 5% sucrose. After removing detached flagella, cells were lysed 1 h at 4 °C in lysis buffer containing 1 mM HEPES (pH 7), 0.5 mM MgCl2, 1% NP-40, and 5000 units DNase. Cell debris was removed by initial centrifugation at 600 x g for 10 min and basal bodies were further purified and concentrated in successive sucrose sedimentation. First, basal bodies were centrifuged in 60%-sucrose cushion at 10,000 x g for 30 min at 4 °C followed by a centrifugation in 40%-70% sucrose gradient at 68,320 x g for 1 h 15 min at 4 °C. Isolated basal bodies were collected at 70% sucrose interface.

### Cloning

A codon-optimized version of a portion of *ccdc81*(1225-1761) gene was synthesized by GeneArt (life technologies) using the codon usage of *Escherichia coli*. It was subsequently cloned in a modified pGEX-6-P vector using the SpeI restriction site. One clone in a good orientation was selected and sequence verified and used to express the corresponding fusion protein. Moreover, Cr*ccdc81*(1225-1761) was also cloned in pmal-c2E vector in a similar fashion.

Similarly, codon-optimized versions of *Crccdc81*-full-length and *CrPOC11* full length were synthesized by GeneArt (life technologies) using the codon usage of *Escherichia coli*. CrCCDC81 construct was subsequently cloned in both modified pGEX-6-P and pmal-c2e vectors and CrPOC11 construct was subsequently cloned in pET151/D-TOPO vector using the SpeI restriction site.

Constructs for expression in U2OS cells were generated in pEGFP-C2 and mCherry-C1 backbone. pEGFP-C2 was digested with EcoRI and KpnI to release the original insert and the backbone was gel purified. The coding sequences of full length CrCCDC81 and CrPOC11 and of the indicated fragments were amplified by PCR from the codon-optimized genes used for recombinant protein expression, using primers carrying an EcoRI site at the 5’ end and a stop codon followed by a KpnI site at the 3’ end. PCR products were digested with EcoRI and KpnI, and ligated into the linearized backbone. Several clones were selected for sequence verification.

### Proteins expression and antibody production

The *Escherichia coli* strain BL21(DE3) (Stratagene) was used for protein expression. In general, bacteria were grown to an OD600 of 0.5 in LB medium at 37°C containing ampicillin. The temperature was switched to 20°C and the culture was induced by the addition of isopropyl 1-thio-β-galactopyranoside (IPTG) to a final concentration of 0.5-1 mM. The protein expression was performed at 20°C for 16h.

The fusion protein GST-CrCCDC81(409-587) was purified using glutathione sepharose beads (GE healthcare). Briefly, the bacterial cell pellet was first resuspended in lysis buffer (50mM Tris pH 7.5, 300mM NaCl, 1mM DTT, proteases inhibitor cocktail, 5mM EDTA) and then incubated with lysozyme and DNase (supplemented with MgSO4 10mM) for 10 minutes on ice. After sonication, 0.5% of Triton was added to the lysate prior centrifugation (12000g for 30 minutes). The obtained soluble lysate was incubated with glutathione sepharose beads for 1h at 4°C on a rotating wheel.

Beads were first washed with lysis buffer supplemented to 1M NaCl and 1% Triton and subsequently washed with lysis buffer. Elution of GST-CCDC81 was performed using 50mM glutathione. The proteins were dialysed overnight against PBS/5% glycerol. GST-CrCCDC81 was cleaved using precision protease (200U for 20mg of protein) for 2hrs at 4°C. The cleaved protein was then subjected to gel filtration using a S200 16/600 column to separate GST from CrCCDC81(409-587). The purified protein CrCCDC81(409-587) was used to immunize one rabbit (Eurogentec). To affinity purify antibodies against CrCCDC81, MBP-CrCCDC81(409-587) was expressed and purified in a similar way but using amylose beads (New England Biolabs). Elution was done using 20mM maltose. Antibodies against CrCCDC81 were column-purified against the MBP-CrCCDC81(409-587) and dialysed against PBS/10% glycerol.

MBP-CrCCDC81-FL overnight culture was grown in LB medium supplemented with 2% glucose at 37°C and culture was induced with 1 mM IPTG at 37°C for 3 hours. The protein was purified as described above.

CrPOC11 antibodies were purified with strip purification protocol. Briefly, 0.5 mg protein was migrated on SDS-PAGE gel and transferred to nitrocellulose, then stained with ponceau S. The position of the protein was marked and cut out. After washing and blocking steps, the strip was incubated in 10-15 mL crude serum for 2 h at room temperature or o/n at 4 °C. After incubation, the serum was removed and the strip was washed once with PBS-Tween and three times with PBS. To elute the antibodies, the strip was placed on a sterile parafilm, protein side facing up and 1 mL of 100 mM glycine pH 2.8-2.5 was applied for 3 mins and the antibodies were collected by carefully pipetting up and down. The eluate was immediately neutralized with 100 µL 1 M Tris pH 8 per mL. The antibody eluate was stored in 50% glycerol at -20 °C.

### CIB1 cassette insertion mapping in CLIP *ccdc81 mutant 1*

Genomic DNA of isolated single clones for *ccdc81* mutant m1 as well as the wild-type strain was performed using Chelex. Briefly, a solution of 5% Chelex resin was prepared in ddH2O (Chelex 100-grade resin, BIORAD). Each single colony from a plate was first resuspended in 20 μL ddH2O, followed by the addition of 20 μL 100% EtOH and 200 μL Chelex 5%. The mixture was vortexed and placed at 100°C for 8 min, then at 4°C for one minute. The samples were spun down at 16 000 g for five minutes in a tabletop centrifuge. Supernatants were collected and contained the extracted genomic DNA. 1 μL of the supernatant was used to perform subsequent PCR reactions. Sets of primers flanking the region of the putative CIB1 insertion site were chosen according to the *Chlamydomonas* library website (https://www.chlamylibrary.org/). To map the cassette-genome junctions and the orientation of the cassette, we used the following primers in the CIB1 cassette: Cassette (1) and Cassette (2).

### Sequence alignment and conservation analysis

Protein sequences of CCDC81 and POC11 homologs were retrieved from UniProt (accession numbers are listed in Table S2). For *C. reinhardtii* CCDC81 and POC11, the sequences used were retrieved from Phytozome. All residue numberings reported in this study refers to those sequences. Sequences were aligned with Clustal Omega using default parameters (Madeira *et al*., 2024). Alignments were visualized in Jalview (2.11.2.7) (Waterhouse *et al*., 2009) and colored with the Clustal Omega color scheme. Per-column conservation scores were calculated in Jalview on a 0-11 scale according to (Livingstone and Barton, 1993), exported, and are reported with respect to the *C. reinhardtii* sequence numbering. Alignments were exported as vector graphics and assembled in Adobe Illustrator.

To display conservation on the predicted structures, the 0-11 conservation scores were grouped into four classes (11, 8-10, 5-7, 0-4) and mapped onto the corresponding residues in ChimeraX, colored purple, dark blue, light blue and grey, respectively.

### Structure prediction and analysis

Structures were predicted with Alphafold3 (Alphafold Server (Abramson *et al*., 2024)) using default settings, and the top ranked model was used throughout. Models were displayed in ChimeraX (1.6.1) (Meng *et al*., 2023) and colored by pLDDT using the standard Alphafold confidence bands (<50, 50-70, 70-90 and >90). Predicted aligned error (PAE) matrices were taken from the Alphafold output and plotted in Python using matplotlib (Hunter, 2007), with dark green indicating low expected position error. For two-chain predictions, interface confidence is reported as the ipTM value from the Alphafold summary confidence output. Models were superimposed in ChimeraX using matchmaker with default parameters, and RMSD values are reported.

### *Chlamydomonas* Immunofluorescence

*Chlamydomonas* cells were deposited on poly-D-lysine coated 12 mm coverslips and incubated for 10 min, protected from light. Coverslips were then fixed for 7 min in – 20°C methanol, washed in PBS, incubated 60 min at room temperature with primary antibodies in 1% bovine serum albumin and 0.05% Triton X-100, washed 15 min in PBS, and incubated 45 min at room temperature with secondary antibodies. Primary antibodies were 1:500 CCDC81 (this study), 1:500 POC11 (this study) 1:400 mouse α-tubulin (ABCD antibodies, aa345), 1:400 β-tubulin (ABCD antibodies, aa344). Secondary antibodies were 1:1000 goat anti-rabbit coupled to Alexa 488 and 1:1000 goat anti-mouse coupled to Alexa 568. Imaging was done on a Stellaris 8 FALCON confocal microscope using an oil immersion 63x objective with a numerical aperture NA of 1.4.

### Human cell lines and cell culture

Human bone osteosarcoma U2OS cells (ATCC-HTB-96) and retinal pigmental epithelial cells hTERT-RPE1 (ATCC-CRL-4000) were grown in Dulbecco’s modified Eagle’s medium supplemented with GlutaMAX, 10% fetal calf serum and penicillin and streptomycin (100 µg/ml) at 37°C in a humidified 5% CO_2_ incubator. Cells were regularly tested for mycoplasma contamination.

### Ultrastructure expansion microscopy (U-ExM)

In U-ExM and iU-ExM experiments, following reagents were used: formaldehyde (FA, 36.5-38%, F8775, SIGMA), acrylamide (AA, 40%, A4058, SIGMA), N, N-methylenbisacrylamide (BIS, 2%, M1533, SIGMA), sodium acrylate (SA, 97–99%, 408220, SIGMA and 7446-81-3, AK Scientific), ammonium persulfate (APS, 17874, ThermoFisher), tetramethylethylendiamine (TEMED, 17919, ThermoFisher), nuclease-free water (AM9937, Ambion-ThermoFisher), poly-D-lysine (A3890401, Gibco) and imaging spacer (iSpacer, 16 mm inner diameter, 22 mm outer diameter, 0.5 mm thickness; SUNJin Lab). Expansion of the cells was performed as previously described (Gambarotto *et al*., 2019; Klena *et al*., 2023) Briefly, *Chlamydomonas* cells deposited on poly-D-lysine coated 12 mm coverslips or human cells that were grown on 12 mm coverslips were incubated for 3 h in an anchoring solution containing 2% AA + 1.4% FA diluted in 1X PBS at 37°C. Next, gelation was performed using monomer solution (10% AA, 19% SA, and 0.1% BIS in 1X PBS) supplemented with 0.5% APS and TEMED by initial incubation of the sample on ice for 5 mins followed by 30 mins incubation at 37°C. Gelation was followed by a denaturation step at 95°C for 1 h 30 mins in denaturation buffer (200 mM SDS, 200 mM NaCl, and 50 mM Tris in nuclease-free water, pH 9.0). Gels were washed three times in ddH_2_O, 10 mins each to remove the denaturation buffer and allow expansion. After the full expansion of the gel, a caliper was used to measure the size to calculate the expansion factor by dividing its diameter to that of the coverslip (12 mm).

### Immunolabelling

Gels were placed in PBS for 20 mins to let them shrink. Then they were incubated in a 1.5 mL Eppendorf tube containing 250 µL primary antibody mixture in PBS-BSA 2% for 2 h 30 mins at 37 °C or overnight at 4°C under agitation. Next, three washes with PBS-Tween 0.1% were performed at room temperature. Then, gels were incubated with secondary antibody mixture in PBS-BSA 2% for 2 h at 37°C. Gels were washed three times in PBS-Tween 0.1% and then washed with ddH_2_O three times 20 mins each before imaging.

### Iterative U-ExM (iU-ExM)

Expansion of the cells was performed as previously described (Louvel *et al*., 2023). Briefly, *Chlamydomonas* cells were incubated in anchoring solution (2% AA + 1.4% FA diluted in 1X PBS) for 3 h at 37°C. Then, a gelation chamber was formed by sticking a double-sided circular spacer on a glass coverslip. The coverslip was glued inside the chamber, cells facing upward. Gelation chamber was placed in a humid chamber on ice, and then monomer solution (MS) (10% AA, 19% SA, 0.1% DHEBA, 0.25% APS and TEMED) was added to fill inside the spacer. The chamber was sealed with a 24 mm round coverslip, and gelation was performed by incubating the sample

15 min on ice followed by 45 min incubation at 37°C. Next, the lid of the chamber was removed and the extra gel around the 12 mm coverslip was cut with a razor blade. The sample was carefully removed from the glass coverslip and placed in a 6-well plate with denaturation buffer (200 mM SDS, 200 mM NaCl, 50 mM Tris-BASE; pH 6.8) with agitation until the gel was detached from the coverslip. Next, the gel was transferred into a 1.5 mM Eppendorf tube filled with 1 mL denaturation buffer and incubated for 1 h 30 min at 85°C. After denaturation, the gel was washed three times 30 min each in ddH_2_O before a last overnight wash for a complete expansion. Next day, the expanded gel was cut in 1,5 x 1,5 cm dimensions for the intermediate antibody staining as previously described for U-ExM protocol. Following the immunostaining step, the 6-well plate containing the gel was placed on ice. The ice bucket was placed on a shaker, and the stained gel was incubated 25 min with activated neutral gel (10% AA, 0.05%DHEBA, 0.1% APS and TEMED in ddH_2_O) under agitation, covered with foil to protect from light. After, the gel was placed on a glass coverslip, and excess solution was gently removed with a kimwipe and was covered with 22 x 22 mm coverslip and incubated in a humid chamber for 1 h at 37°C. Next, the neutral gel was incubated in the anchoring solution (2% AA, 1.4% FA) for 3 h on a shaker at 37 °C. The gel was washed in PBS for 30 min and then incubated for 25 min in 2^nd^ monomer solution (10% AA, 19% SA, 0.1% BIS, 0.1% APS and TEMED) on ice placed on a shaker as the previous step. Next, the gel was placed on glass coverslip, and excess monomer solution was gently removed using a kimwipe. Then, the gel was covered with a 22 x 22 mm coverslip and incubated for 45 min in a humid chamber at 37 °C. After final polymerization, the entire gel was incubated in 200 mM NaOH solution for 1 h under agitation at room temperature protected from light, for dissolution of the first and neutral gels. Following the dissolution, the gel was washed thoroughly with PBS before successive water baths to fully expand the sample. The final ddH_2_O was done overnight.

### Imaging

Expanded gels were cut into pieces to fit them in a 36 mm metallic imaging chamber. The excess water was removed from the gels using a kimwipe. After finding the correct orientation of the cells in the gel, it was mounted on a 24 mm round precision coverslip coated with poly-D-lysine to stabilize it. Image acquisition was performed with a 63X 1.4 NA oil immersion objective with an inverted widefield Leica DMi8 Thunder microscope or with a confocal Leica Stellaris 8 FALCON microscope. The microscope parameters were controlled using the LAS X software. Denoising for widefield images were performed by selecting “small volume clearing” mode on Thunder and water as the “mounting medium” and 3D stacks were acquired with a voxel size of 0.1013 × 0.1013 × 0.2131 μm. Confocal images were processed with lightning mode at max resolution, adaptive as “strategy” and water as the “mounting medium” to generate deconvolved images with adapted voxel size.

### Antibodies used in this study

The primary antibodies used in this study are as follows: alpha-tubulin (aa345, ABCD antibodies; 1:250), beta-tubulin (aa344, ABCD antibodies; 1:250), CrCCDC81 (this study, 1:250), POC11 (this study, 1:250), CCDC81 Ab#1 (PA5-48210, ThermoFisherSci, 1:200), CCDC81 Ab#2 (ab126514, Abcam, 1:200), CCDC81 Ab#3 (orb513188, Biorbyt, 1:200), CCDC81 Ab#4 (orb423095, Biorbyt, 1:200), CCDC77 (26369-1-AP, Proteintech, 1:250), WDR67 (HPA023710, Atlas antibodies, 1:250), Acetylated tubulin (T7451, Merck Sigma-Aldrich, 1:500 for IF), POC5 (A303-341A, Bethyl, 1:250), CEP63 (16268-1-AP, Proteintech, 1:500), C2CD3 (HPA038552, Atlas antibodies, 1:250), CEP164 (22227-1-AP, Proteintech, 1:250), Centrin (clone 20H5, 04-1624, Merck Millipore, 1:500 for IF). The secondary fluorescent antibodies are from Invitrogen (anti-rabbit 488 - A11008, anti-rabbit 568 – A11036, anti-mouse 488 A11029, anti-mouse 568 – A11004, anti-guinea pig Cy5 – Jackson Immuno Research 706-175-148) and used at 1:1000 dilution for IF and 1:500 dilution for U-ExM.

### Immunofluorescence microscopy

U2OS and RPE1 cells grown on coverslips were fixed in methanol at -20°C for 3 min and washed three times with PBS. Cells were incubated with primary antibodies diluted in PBS with 2% BSA for 1h at room temperature, washed three times with PBS containing 0.1% Tween-20, and incubated with secondary antibodies for 1 h at room temperature. Microtubules were stained with a mixture of α- and β-tubulin antibodies, centrioles with anti-centrin-2, primary cilia with acetylated tubulin and CCDC81 with Ab#1 to Ab#4 (antibodies and dilutions are listed above). After washing, coverslips were mounted in glycerol mounting medium containing DAPI and DABCO (Abcam, ab188804). Cell cycle stages were assigned from centriole number and DNA morphology. Images were acquired on a Leica Thunder widefield microscope with a 63X 1.4 NA oil objective and processed with ImageJ.

### Microtubule pelleting assay

25 µg of tubulin (5µl tubulin at 5mg/ml) was first diluted in 20 µl of BRB80 buffer (80 mM K-PIPES, 1mM EGTA, 1 mM MgCl2) supplemented with 1 mM GTP and 1mM DTT and incubated for 5 min at 4°C. Next, tubulin was polymerized stepwise by the first incubation at 37°C for 10 min, followed by the addition of 0.1 µM (5 min), 1 µM (5 min), and 10 µM taxol (10 min) at 37°C. Following microtubule polymerization, 20 µg of recombinant protein, either MBP-CrCCDC81-FL or His-POC11-FL, was incubated in BRB80 buffer for 30 min at RT. The sample mixture was then loaded on a glycerol cushion and centrifuged in a TLA-55 at 50’000 rpm for 30 min at 37°C. The supernatant was collected, while the pellet was resuspended in SDS sample buffer. 50% of total supernatant and pellet were then loaded on a 10% SDS-PAGE gel, which was then stained using Coomassie.

### Microtubule binding assay

Microtubule binding assay was performed as described previously (Tao *et al*., 2016). Briefly, 20 µg tubulin (4 µl tubulin at 5 mg/ml) was diluted in total volume of 20 µl of BRB80 buffer to reach 10 µM concentration (1 mg/ml). Microtubules were polymerized as described above using increasing taxol concentrations. After the centrifugation, the pellet was gently washed with 0.5% Triton-X and resuspended in 20 µl warm BRB80 buffer to reach 10 µM microtubule concentration. 30 µl reaction was set in Buffer T (20 mM Tris pH 8.0, 150 mM KCl, 2 mM MgCl2, 1 mM DTT, protease inhibitors) with 6 µl microtubules (2 µM final concentration), 20 µM taxol and 100 nM or 500 nM of MBP, MBP-CCDC81 or His-POC11 for 30 min at room temperature. Poly-D-lysine coated coverslips were washed with 100% EtOH and dried. Coverslips were placed in a 24-well plate and filled with 150 µl 20% glycerol cushion. The reaction was added on the glycerol cushion and spun at max speed (3428 g, Centrifuge 5920 R). Coverslips were fixed with MeOH and stained for tubulin and MBP, CrCCDC81 or POC11. For copolymerization assays, equimolar concentrations of MBP-CCDC81 or His-POC11 were mixed with tubulin, and polymerization was performed as described.

Samples were imaged with Leica SP8 inverted confocal microscope (**Fig 3**; **Fig S5**) or with TIRF module of Leica DMi8 widefield microscope (**Fig S5**).

### TIRF microscopy

Porcine brain tubulin, biotin-labeled tubulin as well as rhodamine and HiLyte-488-labeled tubulin were purchased from Cytoskeleton Inc. (T240, T333P, TL590M and TL670M, respectively).

*GMPCPP-stabilized biotin-labelled microtubules* were polymerized from 5μl of 4 mg/ml rhodamine-labelled biotinylated porcine-brain tubulin (12% biotin tubulin, 20% rhodamine-labeled tubulin, 68% unlabelled tubulin) for 1 hour at 37°C in BRB80 buffer (80 mM K - PIPES pH 6.8, 1 mM K - EGTA, 2 mM MgCl2) supplemented with 2.7 mM GMPCPP (Jena Bioscience, NU-405). The polymerized microtubules were centrifuged for 30 min at 18000 x g in a microfuge (Eppendorf, Centrifuge 5424 R). After centrifugation the pellet was resuspended in BRB80 at RT.

Total internal reflection fluorescence (TIRF) microscopy experiments were performed on an inverted widefield Leica DMi8 Thunder microscope equipped with TIRF module, 100x NA 1.4 oil immersion objectives and with Leica K5 DOC camera. Rhodamine-labelled biotinylated microtubules and HiLyte488 labelled tubulin were visualized sequentially by switching between microscope filter cubes for GFP and mCherry channels. The imaging setup was controlled by LasX software (Leica).

Flow chambers for TIRF imaging assays were prepared as described previously (Fink *et al*., 2009). Coverslips of two different sizes: 22x22 mm and 18x18 mm were cleaned using air plasma (Hirst *et al*., 2020) by hexamethyldisilazane (HMDS, 440191-100ML, Sigma-Aldrich). Chambers were constructed by five thin strips of parafilm M (VWR) (23 mm long, 2 mm width) which were placed on to larger coverslip and the smaller coverslip was placed on top of parafilm strips, forming a coverslip “sandwich”. The assembled structure was transferred onto a heating plate at 60°C, and gentle pressure was applied to the top of the coverslip using tweezers to create firmly sealed flow channels. Subsequently, the assembled flow chambers were placed onto a custom-built microscopy holder, ensuring close contact between the chamber and the microscope objective from below for optimal imaging conditions. Channels were treated with 6 ug/ml anti-biotin solution (Sigma, B3640, 1 mg/ml in BRB80) after 5 minutes incubation, followed by one-hour incubation with 1% Pluronic F127 diluted in BRB80 (Sigma, P2443).

*Microtubule dynamic assay.* Rhodamine-labelled biotinylated microtubules were immobilized and subsequently flushed in the assay buffer (BRB80, 1 mM GTP, 0.2% Tween20, 0.5 mg/ml Casein, 20 mM D-glucose, 0.22 mg/ml glucose oxidase and 20 ug/ml catalase) with free 16 μM HiLyte488-tubulin (25% HiLyte488-tubulin, 75% unlabelled tubulin) in the absence or presence of 100 nM CCDC81 or 1 μM CCDC81. Time-lapse image sequences were recorded for 15 min at the rate of 1 frame per 10 s with an exposure time 150 ms. Data from three independent experiments was collected, each experiment was repeated on different days.

*Image analysis.* Microscopy data were analyzed using ImageJ (FIJI) (Schindelin *et al*., 2012). Microtubules dynamic instability parameters as polymerization rate, depolymerization rate, frequency of catastrophes and frequency of rescues were analysed using Kymographs, which were generated using KymographBuilder plug-in in FIJI by manually fitting lines along the microtubules.

### In vitro pull-down assay

Pulldowns were performed with purified proteins at 4°C with constant rotation. For pulldowns with His-POC11 as bait, 20 µg of full-length His-POC11 was incubated with 20 µl of a 50% slurry cobalt resin (HisPur Cobalt Resin, Thermo Scientific, #89964) in 150 µl PBS for 1 h. Beads were collected by centrifugation (500 x g, 3 min, 4 °C), a sample of unbound fraction was kept and the beads were washed twice with 1 ml PBS supplemented with 150 mM NaCl. The washed beads were incubated with 20 µg of full-length MBP-CrCCDC81 in 150 µl PBS for 2 h, then collected and washed as above. Bound proteins were eluted by boiling the beads in 40 µl SDS sample buffer. As a negative control, 20 µg purified MBP was incubated with His-POC11 immobilized beads in parallel. For the reciprocal pulldown, 20 µg of full-length MBP-CrCCDC81 was immobilized on 50 µl amylose resin (New England Biolabs, E8021S) and incubated with 20 µg His-POC11 under the same conditions.

To test tubulin interaction, His-POC11 or MBP-CrCCDC81 bound beads were incubated with 20 µg purified tubulin (Cytoskeleton Inc., T240) and processed as described above. To test whether three proteins associate together, His-POC11 bound beads were first incubated with MBP-CrCCDC81 and then with tubulin.

Inputs, unbound fractions and eluates were separated by SDS-PAGE on 4-20% gradient gels (BioRad, #4561094) and stained with One-Step Blue (Biotium, #21003-1L).

### Displacement assay

U2OS cells grown on coverslips were transfected at 80% confluency in a six-well plate using jetPRIME reagent (Polyplus-transfection) following the manufacturer’s instructions, with 2.5 µg of total DNA. To map the microtubule-binding regions, cells were transfected with GFP-tagged full-length CrCCDC81 or CrPOC11, or with GFP-tagged fragments of CrCCDC81 (residues 1-170, 1-200, 90-200, 1-250, 250-480, 480-800 and 200-800) or CrPOC11 (residues 1-150, 150-300, 300-480, 1-388, 251-450, 300-450 and 391-481). For displacement assays, mCherry-tagged full-length CrCCDC81 was co-expressed with GFP-tagged full-length (residues 1-480), N-Terminal (residues 1-150), middle (residues 150-300) or C-terminal (residues 300-480) fragments of CrPOC11. Similarly, mCherry-tagged full-length CrPOC11 was co-expressed with GFP-tagged middle (residues 250-480) or C-terminal (residues 300-480) fragments of CrCCDC81, using equal amounts of each plasmid. The medium was changed 4-6 h after transfection and the proteins were expressed for 24 h before cells were fixed and stained for tubulin as described in immunofluorescence microscopy.

*Cold-stability assay.* 24 h after transfection, untransfected cells and cells expressing GFP-CrCCDC81 or GFP-POC11 were placed on ice for 10 min, fixed immediately in cold methanol and stained as above. Cells were scored as having cold-stable microtubules when cytoplasmic microtubule network remained visible. For displacement assays, cells expressing both constructs were scored for the presence of the GFP-tagged fragment on microtubules.

### siRNA-mediated protein depletion

RPE-1 cells were plated in 6-well plates at 100,000 cells per well for the control siRNA and at 150,000 cells per well for the siRNAs against CCDC81 and CCDC77 and transfected 6 h later using Lipofectamine RNAiMAX (Invitrogen) according to the manufacturer’s instructions. All siRNAs were Silencer Select (Thermo Fisher) and were used at 20 nM total concentration: negative control siRNA (#4390843), siCCDC77 (s38908; sense 5’-GACGUAUCCUGGAAGUAGAtt-3’), or a pool of two pre-designed siRNAs against CCDC81 at 10 nM each (s34106: sense 5’-GAGCUGGAGCGAGUAAAUAtt-3’, antisense 5’-GAGCUGGAGCGAGUAAAUAtt-3’, antisense 5’-UAUUUACUCGCUCCAGCUCag-3’; s34107: 5’-GUCCUGCGCUUAAUGCUCUtt-3’, antisense 5’-AGAGCAUUAAGCGCAGGACtg-3’). For the double depletion, cells were transfected with 5 nM of each CCDC81 siRNA and 10 nM of siCCDC77. The medium was changed after 5-6 h after transfection, and a second transfection was done 48 h after the first one without changing the medium. Cells were analyzed 96 h after the first transfection and were serum-starved for the last 24 h before fixation to induce ciliogenesis. Cells were fixed and stained for acetylated tubulin as described in Immunofluorescence. The percentage of ciliated cells was scored, and cilia length was measured from the acetylated tubulin signal. For centriole analysis, depleted cells were processed as described in U-ExM. Knockdown efficiency was assessed from the centriolar CCDC77 and CCDC81 signals in U-ExM.

### Cryo-electron microscopy

#### Sample preparation and vitrification

Taxol-stabilized microtubules were prepared as described for the binding assays, using 2 µl of tubulin in a final volume of 40 µl BRB80. Quantifoil R2/1 Cu 300-mesh holey carbon grids (Quantifoil Micro Tools) were glow discharged for 60 s at 30 mA (negative polarity) in a GloQube (Quorum Technologies). Grids were prepared in a Leica EM GP2 (Leica Microsystems) at 37°C and 90% relative humidity. 5 µl microtubules were applied to the grid, incubated for 30 s and back-blotted 1 s. MBP-CrCCDC81 (5 µl at 1.57 µM; purified as described above) was then applied, incubated for 30 s and back-blotted for 1 s, and the grid was plunge-frozen in liquid ethane. Undecorated microtubules were prepared in the same way, without the MBP-CrCCDC81 step.

#### Data collection

Micrographs were recorded in one screening session on a Talos Arctica (Thermo Fisher Scientific) operated at 200 kV at the Dubochet Center for Imaging in Geneva. The microscope had a Falcon 4i detector and a Selectris X energy filter (10 eV slit width). Single 9.2 s exposures (not dose-fractionated) were recorded in linear mode at a nominal magnification of 130,000x (0,9024 Å/pixel) and a nominal defocus of -3.0 µm, with a total exposure of 20-32 e^-^/Å^2^ per image. Five micrographs of undecorated microtubules and 22 of microtubules incubated with MBP-CrCCDC81 were analyzed.

#### Single particle processing

The 22 micrographs of MBP-CrCCDC81 incubated microtubules were processed in CryoSPARC v4.7.1 (Punjani *et al*., 2017) following the MiCSPARC protocol (Zhang *et al*., 2026). They were single exposures, so no motion correction was needed, and CTF parameters were estimated with Patch CTF. Microtubules were traced with the filament tracer. Segments were extracted in 567 Å boxes and cleaned by 2D classification. Picks were then extended along each filament with MiCSPARC’s filament extrapolation (2,398 segments). Segments were then re-extracted at 1.97 Å/pixel (628-pixel boxes Fourier-cropped to 288 pixels) and classified in 2D again, leaving 2,178 segments.

Protofilament numbers were assigned following MiCSPARC. Helical refinements were run with the theoretical parameters of the 11-3 to 16-4 lattices, and the best-resolved one (13-3) was used to generate references by heterogeneous refinement, followed by per-filament consensus assignment. Of the segments, 58% came from 13-, 23% from 12-, 17% from 11-, and 3% from 14-protofilament microtubules. Only the 12- and 13-protofilament segments were processed further, as two separate sets. Each set was refined with the theoretical helical parameters of its lattice (12-3 or 13-3), and then without imposed symmetry. CTF refinement was skipped because the helical maps were limited to about 10 Å. In-plane and azimuthal angles were made consistent along each filament with MiCSPARC’s psi and phi unification scripts, each followed by local refinement. Segments were then symmetry-expanded 12- or 13-fold with the refined helical parameters and locally refined.

One protofilament was isolated with a wedge-shaped mask. The rest of the microtubule was removed by particle subtraction, and the protofilament was locally refined in 144-pixel boxes (1.97 Å/pixel). The two sets were merged (21,105 protofilament particles) and locally refined to 6.7 Å resolution. These particles were sorted into four classes by 3D classification, and each class was reconstructed separately (class 0: 5,186 particles, 7.6 Å). Class 0 was locally refined, local resolution was estimated, and maps were locally filtered in CryoSPARC. Maps were displayed in UCSF ChimeraX (Meng *et al*., 2023).

### Quantification and Statistical analysis

#### Image analysis and representation

Images were analyzed using Fiji/ImageJ (Schindelin *et al*., 2012). All intensity measurements were performed on raw data. Deconvolved images were used only for representation. Brightness and contrast were adjusted linearly and applied identically to all conditions being compared within a figure panel.

#### Expansion factor

For every U-ExM and iU-ExM experiment, the diameter of the expanded gel was measured with a caliper, and the expansion factor was calculated as the ratio of the gel diameter to that of original coverslip (12 mm). The expansion factor of the corresponding gel was applied to all measurements before plotting, so all distances reported in the text and in the graphs correspond to pre-expansion values. Scale bars in the images of expanded samples are given in expanded-gel dimensions and are not corrected for expansion factor.

#### Measurements of expanded centrioles

Centrioles were selected based on tubulin signal and assigned to longitudinal or top views. Lengths, relative positions and coverage were measured on side views and diameters on top views as previously described (Laporte *et al*., 2024). Briefly, Fluorescence distributions were obtained with a line scan drawn along the centriole long axis with the plugin “PickCentrioleDim”, and the start and end of each signal were defined as 50% of the maximum peak intensity at the two extremities. Tubulin was used as the reference channel and its proximal coordinate was set to 0 and the same shift was applied to the protein of interest, so that relative distances between the two signals were preserved. Negative values therefore indicate the positions proximal to the start of the tubulin signal. Diameters were measured on top views as the distance between the two intensity maxima of a line scan drawn across the centriole.

#### Scoring of centriole defects

Centriole morphology was scored on U-ExM images of RPE-1 cells stained for tubulin and the indicated markers. A centriole was classified as abnormal when it displayed incomplete or open microtubule walls, loss of the proximal or distal region, deformed architecture, or isolated microtubule blades. Cells were then classified according to whether none, one or both centrioles were abnormal.

#### Depletion efficiency

Depletion was quantified from immunofluorescence images by measuring the mean intensity of the protein of interest in a fixed size region centered on the centrosome/basal bodies and in an identically sized region in the adjacent cytoplasm.

The same region size was used for all images within an experiment type (regular immunofluorescence or U-ExM). The cytoplasmic value was subtracted as background, and centrosomal intensities were normalized to the mean of the control siRNA condition of the same experiment.

#### Statistics

Statistical analyses were performed in GraphPad Prism. Data are presented as mean ±SD unless otherwise stated. Conditions were compared by one-way ANOVA followed by Dunnet’s multiple comparison test when all conditions were compared with a single control, or by two-way ANOVA followed by Tukey’s multiple comparison test when two factors were considered. The test used for each quantification is indicated in the corresponding figure legend. In figure legends, n indicates the number of independent experiments and N the number of centrioles, cells, or microtubules measured. Both numbers are reported for every quantification in figure legends. All experiments were performed at least three times independently unless otherwise stated.

## Data availability

All data are available upon request.

## Author Contributions

U.B. performed and analyzed the experiments of the paper with some initial help from S.B. D. K. performed and analyzed the TIRF experiments. V.H. and P. G. designed and supervised the project. U.B., V.H, and P.G. wrote and revised the manuscript.

## Acknowledgments

We thank Vincent Louvel for the initial iU-ExM experiments and Davide Gambarotto for initial U-ExM staining on isolated *Chlamydomonas* basal bodies, Natacha Olieric and Michel Steinmetz for the purification of the CrPOC11-FL protein. We thank Lorène Bournonville for critical reading of the manuscript. This work was supported by the Swiss National Foundation (SNSF) 310030_205087, awarded to V.H. and P.G; the University of Geneva and the European Research Council ERC Consolidator Grant ISAC, administered in Switzerland by SERI (contract MB22.00075), awarded to P.G.

## Supplementary Figure legends

**Figure S1.**
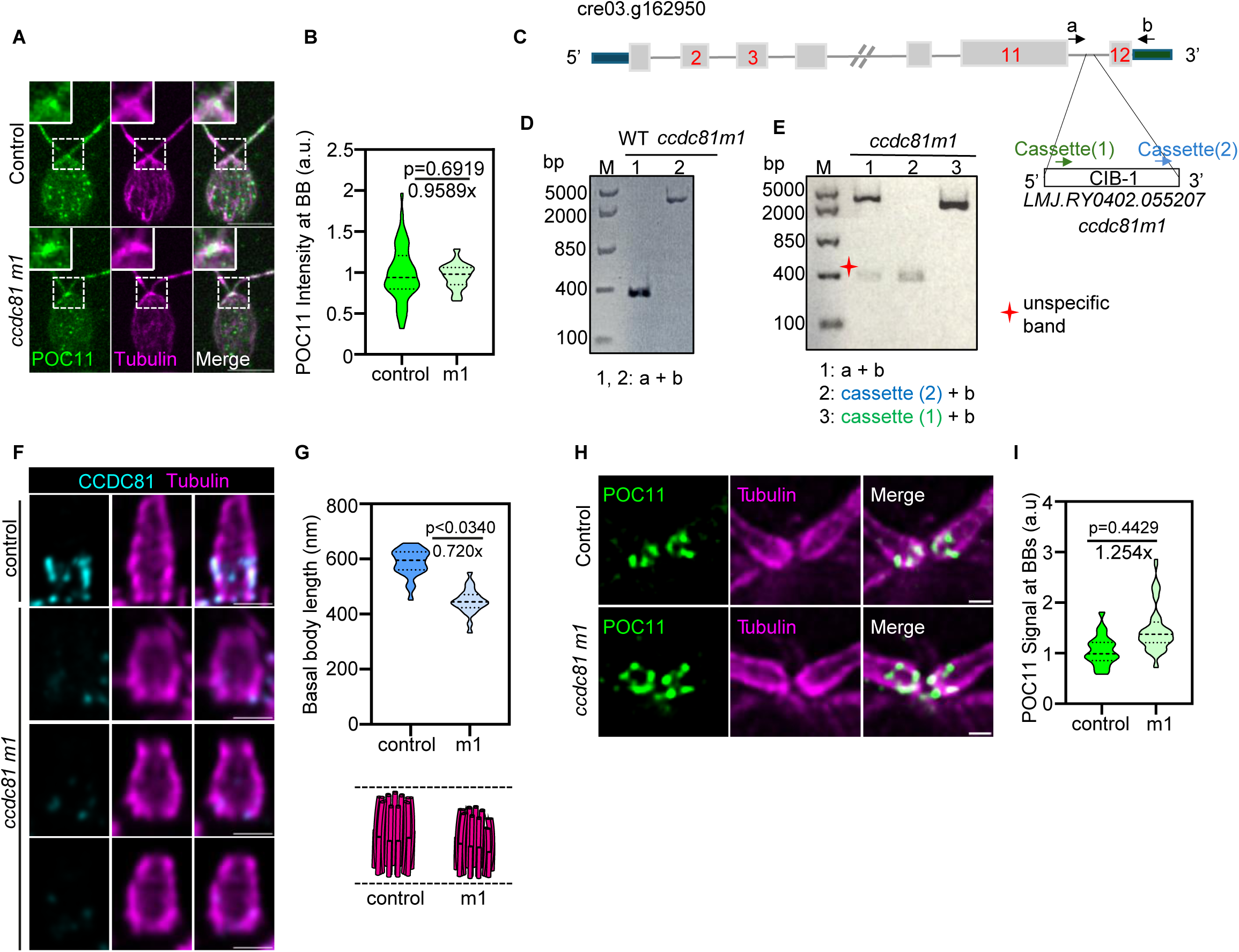
Characterization of the ccdc81 mutant and POC11 localization in *Chlamydomonas*. **(A)** Immunofluorescence microscopy of control *Chlamydomonas* cells (top panel) and *ccdc81 m1* mutant cells (bottom panel) stained with antibodies against POC11 (green) and α/β tubulin (magenta). Dashed white boxes indicate the position of the basal bodies which corresponds to in 4-fold magnified insets. Scale bar = 5 μm. (**B**) Quantification of POC11 normalized signal intensity at basal bodies in control and *ccdc81 m1* mutant cells. Control, *N* = 77, *n* = 2 (mean= 1.000); *ccdc81 m1*, *N* = 117, *n* = 2 (mean= 0.9589). Nested t test, two-tailed. (**C**) Schematic representation of the gene cre03.g162950 (from Phytozome v12.0). The CIB1 cassette insertion site for *ccdc81 m1* is indicated. 5’ and 3’ UTR are shown in blue and green respectively. Exons are in grey and introns as thin black lines. Primers used for mapping are shown as black arrows (a, b). **(D, E)** PCR amplification from either wild-type (WT) or *ccdc81 m1* genomic DNA using primers as indicated. Expected size of WT (a+b)= 344bp (D, lane 1). Note that when a CIB1 cassette is inserted we observed a band>2.5kb (D, lane 2). The orientation of the CIB1 cassette has been determined using either cassette (1) or cassette (2) primers (E). **(F)** Expanded *C. reinhardtii* basal bodies of control (top panel) and *ccdc81 m1* mutant (lower panels) stained for α/β tubulin (magenta) and CCDC81 (cyan), shown in longitudinal and top view. Scale bar = 1 μm. **(G)** The quantification of basal body length in expanded cells. Control, *N* = 67, *n* = 3 (mean = 604.7 nm); ccdc81 m1, N = 53, n = 3 (mean = 435.8 nm). Nested t test, two-tailed. The illustration below summarizes the reduction in basal body length in the mutant. **(H)** Expanded *C. reinhardtii* cells showing the basal bodies of either control (top panel) or *ccdc81 m1* (bottom panel) stained for α/β tubulin (magenta) and POC11 (green). Scale bar = 1 μm. **(I)** The quantification of the POC11 normalized signal intensity at expanded basal bodies in control and CCDC81 mutant cells. Control, *N* = 99, *n* = 3 (mean= 1.0); CCDC81 m1, *N* = 131, *n* = 3 (mean= 1.25). Nested t test, two-tailed.

**Figure S2.**
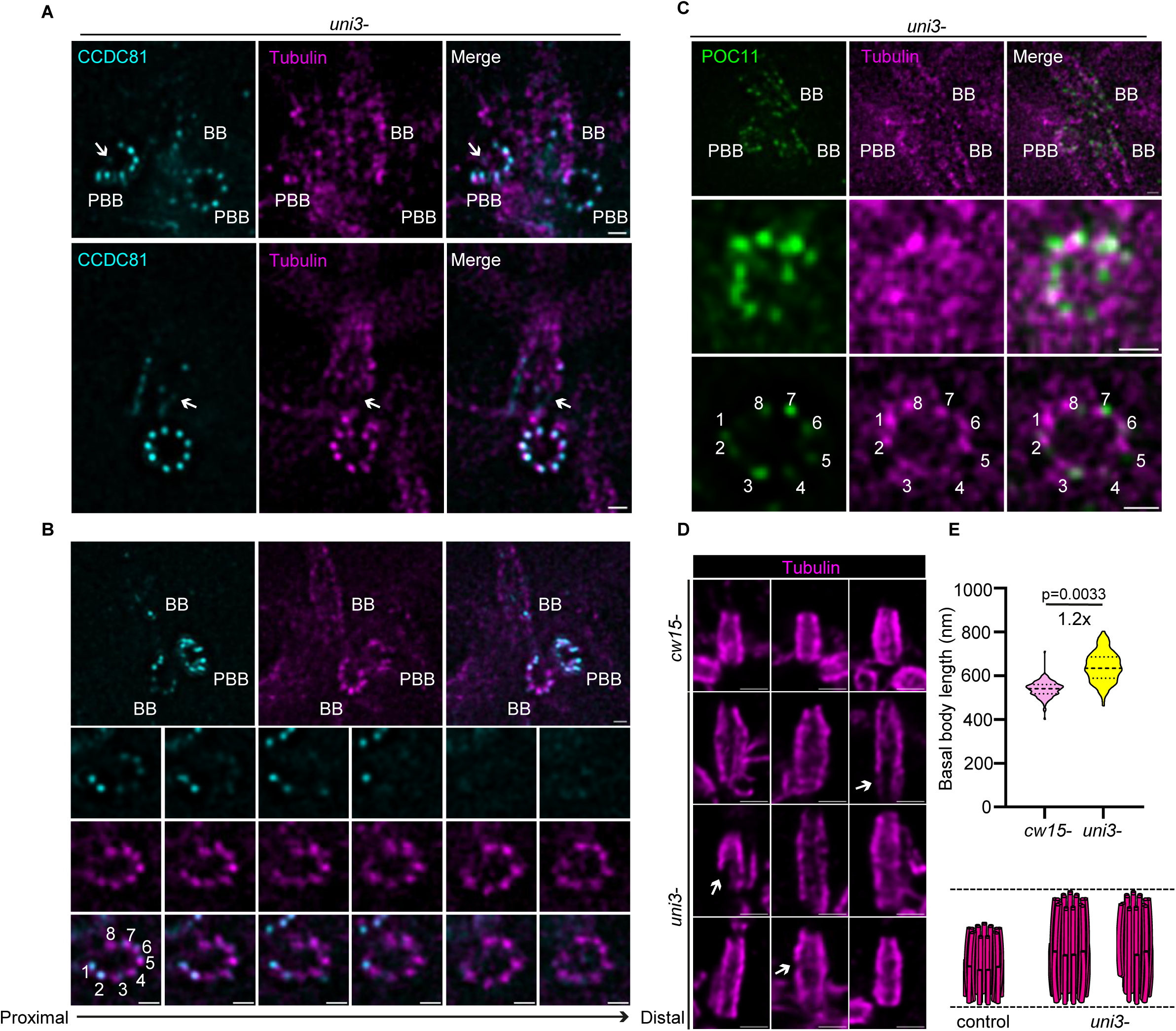
Loss of δ-tubulin disrupts basal-body architecture in *Chlamydomonas*. N, measurements; n, independent experiments. Scale bars indicate pre-expansion dimensions. **(A)** iU-ExM of *C. reinhardtii* δ-tubulin mutant (*uni3-*) cells stained for α/β tubulin (magenta) and CCDC81 (cyan). Two different cells are shown, in lateral view (top panel) and top view (bottom panel). White arrows point to the missing tubulin signal. Basal bodies (BB) and pro-basal bodies (PBB) are marked. Scale bar = 1 μm. **(B)** A further *uni3-* basal body shown from the top view with missing tubulin signal. Successive z-stacks running from proximal to distal, oriented by the CCDC81 staining, show that the missing tubulin signal is in the proximal part. Microtubule triplets (MTTs) are numbered 1 to 8; the ninth triplet is absent. Scale bar = 1 μm. **(C)** iU-ExM of *uni3-* cells stained for α/β tubulin (magenta) and POC11 (green), shown as an overview (top panel), a magnified view (middle panel) and the top view (bottom panel). The staining shows disorganized MTTs and a missing triplet. Scale bar = 1 μm. **(D)** Expanded *C. reinhardtii* basal bodies of *cw15-* (top panels) and the δ-tubulin mutant *uni3-* (bottom panels) in longitudinal view, stained for α/β tubulin (magenta). Scale bar = 1 μm. **(E)** The quantification of basal body length in expanded cells. *cw15-*, *N* = 363, *n* = 3 (mean = 536.2 nm); *uni3-*, *N* = 273, *n* = 3 (mean = 645.1 nm). Nested t test, two-tailed. The illustration below summarizes the increase in basal body length in the mutant.

**Figure S3.**
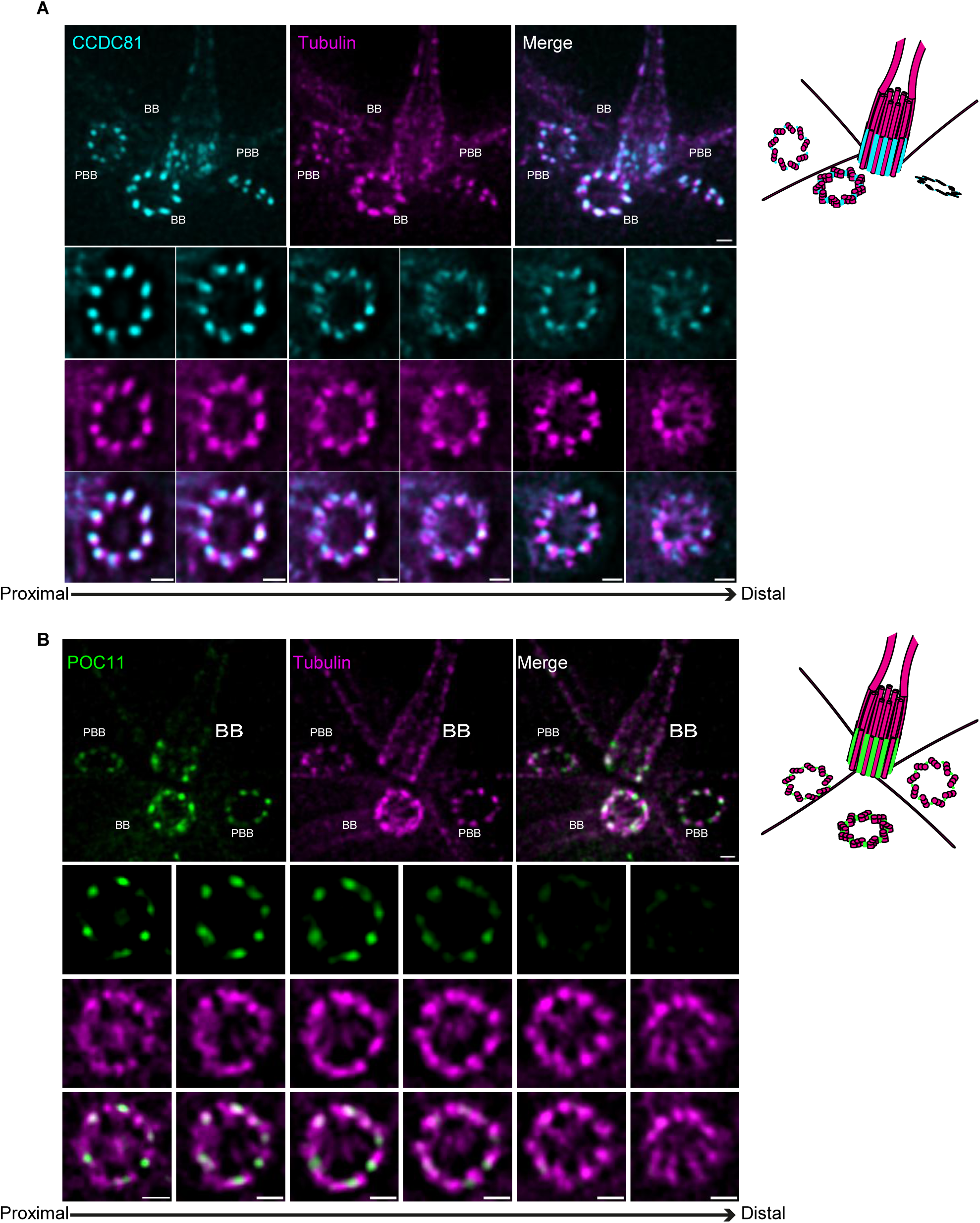
CCDC81 and POC11 are restricted to the proximal basal-body region. (**A, B**) iU-ExM of *Chlamydomonas reinhardtii* cells stained for α/β tubulin (magenta), and CCDC81 (cyan) (A) or POC11 (green) (B). Top panels show the basal body region, where basal bodies (BB) and pro-basal bodies (PBB) are marked. Below, one BB is shown from the top view as successive z-planes going from the proximal to the distal part, as indicated by the arrow. For each cell, the top row shows CCDC81 (A) or POC11 (B), the middle row shows α/β tubulin, and the bottom row shows the merge. The signal of both proteins is present in the proximal planes and is lost towards the distal part, while the tubulin signal is present throughout. Scale bars = 1μm

**Figure S4.**
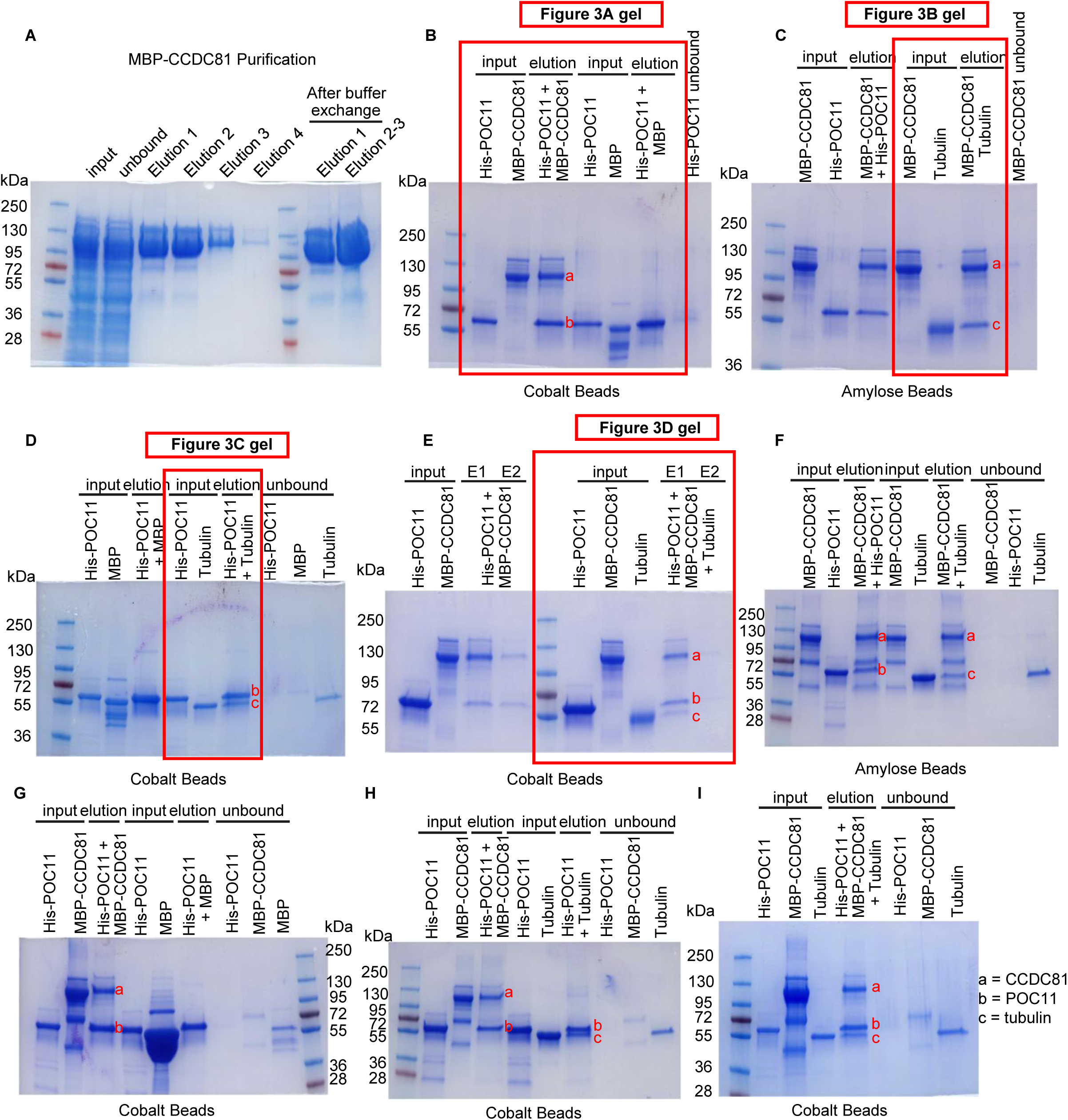
Protein purification, uncropped gels and independent pulldown replicates. Gels are Coomassie-stained; markers are in kDa. Lane identities are as labelled above each gel. Several experiments were run on each gel, so individual replicates are distributed across panels. **(A)** Purification of MBP-CCDC81, showing the input, the unbound fraction and successive elutions, together with the pooled elutions after buffer exchange. **(B–E)** Uncropped gels corresponding to Figure 3A (B), Figure 3B (C), Figure 3C (D) and Figure 3D (E). Red boxes indicate the regions shown in the main figure. Lanes outside the boxes include the unbound fractions and, in (C) and (D), additional pulldowns. **(F–I)** Further independent replicates of the pulldown assays in Figure 3, each performed with an independent protein preparation. The resin used is indicated below each gel. (F) shows the reciprocal pulldown, in which MBP-CCDC81 rather than His-POC11 is immobilized. Across panels **(B–I)**, replicates are distributed as follows: His-POC11 on cobalt resin with MBP-CCDC81, panels B, E, G and I (n = 4); His-POC11 on cobalt resin with MBP alone as a negative control, panels B, D and G (n = 3); MBP-CCDC81 on amylose resin with His-POC11, the reciprocal pulldown, panels C and F (n = 2); His-POC11 on cobalt resin with tubulin, panels D and H (n = 3); MBP-CCDC81 on amylose resin with tubulin, panels C and F (n = 3); and His-POC11, MBP-CCDC81 and tubulin together, panels E and I (n = 4).

**Figure S5.**
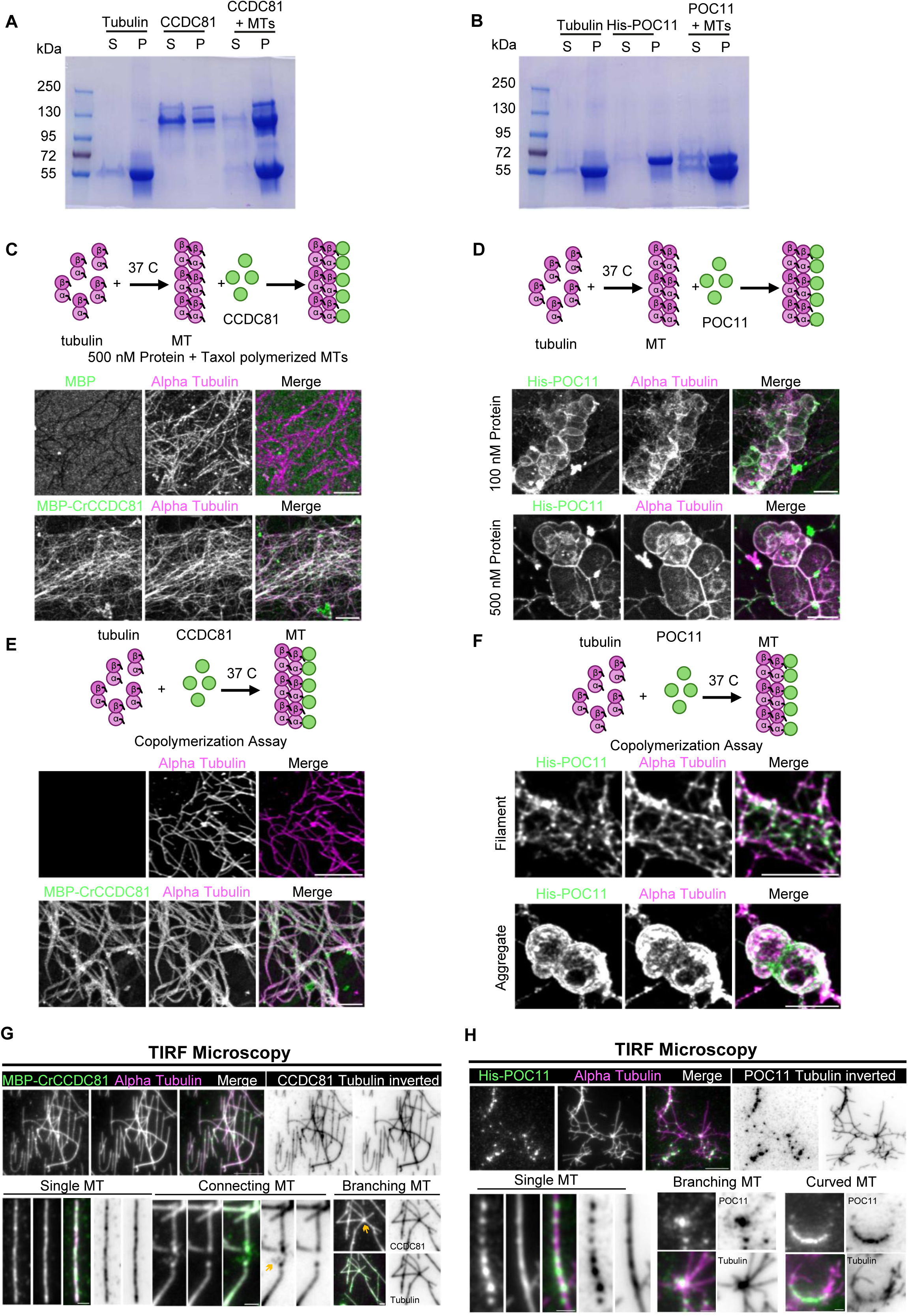
Biochemical and imaging assays of CCDC81 and POC11 association with microtubules. All assays are representative of 3 independent experiments. “Inverted” panels in (D) and (H) show the inverted greyscale of the individual channels. **(A, B)** Microtubule (MT) pelleting assay for CCDC81 (A) and POC11 (B). The soluble (S) and pellet (P) fractions were loaded on an SDS-PAGE gel and stained with Coomassie blue. Molecular weight markers are indicated in kDa. Note that a fraction of CCDC81 pellets without MTs, while all POC11 pellets by itself, making the MT interaction inconclusive from this assay. **(C, D)** MT binding assay. The schemes at the top summarize the experiment. (C) 500 nM MBP alone as a negative control, or MBP-CCDC81; the corresponding 100 nM condition is shown in Figure 3E. (D) His-POC11 at 100 nM and 500 nM. Coverslips were stained for α/β tubulin (magenta) and MBP, CCDC81 or POC11 (green). Scale bars = 5 μm. **(E, F)** MT copolymerization assay. (E) Tubulin alone and MBP-CCDC81. (F) His-POC11, which both decorates filaments (Filament) and forms aggregates (Aggregate). Scale bars = 5 μm. **(G, H)** *In vitro* polymerized MTs were mixed with MBP-CCDC81 (G) or His-POC11 (H), fixed on coverslips and stained for α/β tubulin (magenta) and CCDC81 or POC11 (green), then imaged by TIRF microscopy to resolve individual MTs. Below, examples of single, connecting and branching MTs (G) and of single, branching and curved MTs (H). Yellow arrows in (H) indicate concentrated CCDC81 at MT tips and clusters. Scale bars = 5 μm for the top panels and 1 μm for the bottom panels.

**Figure S6.**
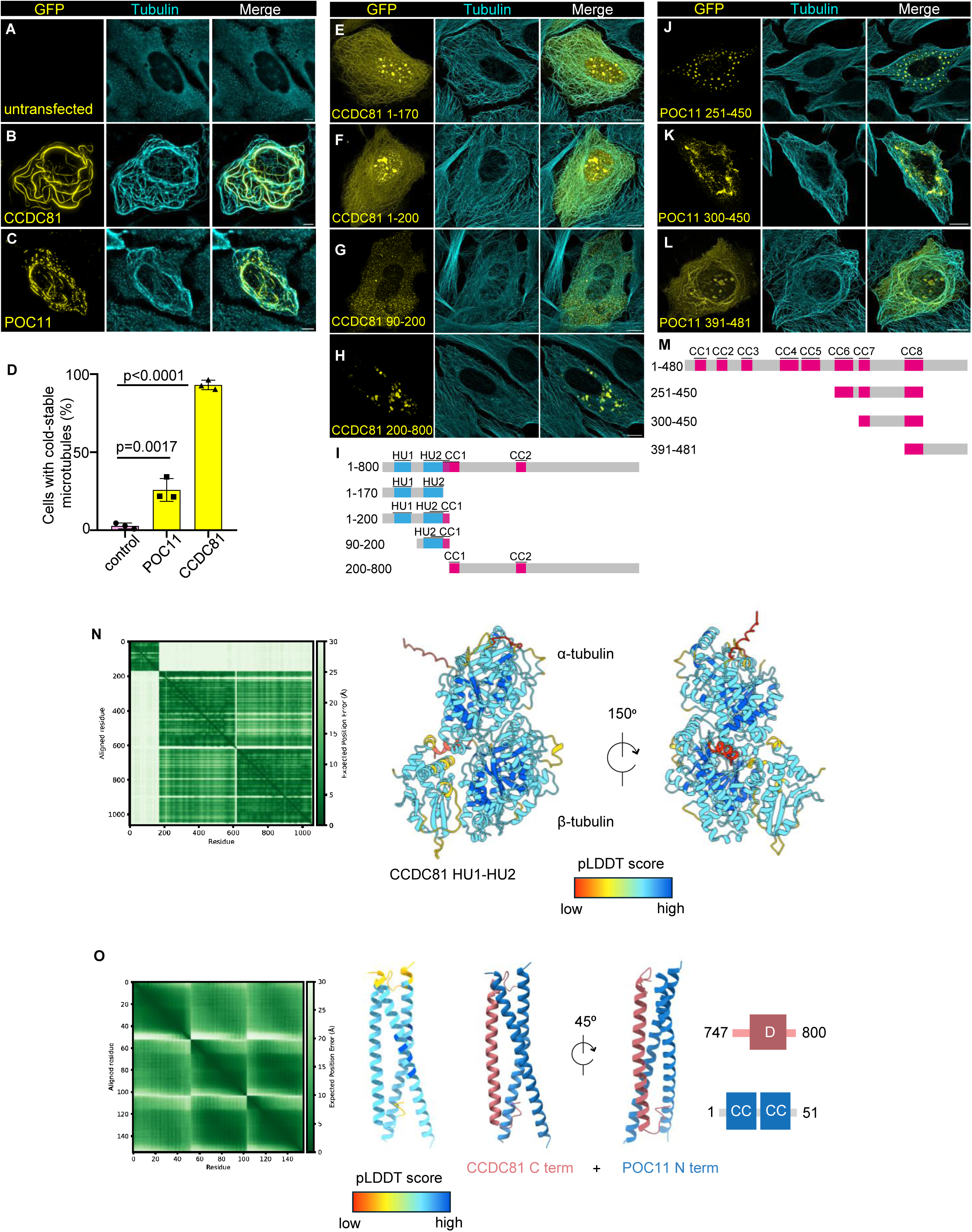
Functional and domain-level characterization of CCDC81 and POC11. Constructs, transfection, fixation and staining were as in Figure 4 unless stated otherwise. **(A–C)** Representative U2OS cells after cold treatment (0 °C, 10 min, on ice): untransfected (A), GFP-CCDC81 (B) and GFP-POC11 (C). Each row shows GFP, α-tubulin and the merge. Scale bars = 5 µm. **(D)** Percentage of cells retaining cold-stable microtubules, scored in untransfected cells and in cells expressing GFP-CCDC81 or GFP-POC11 from the same experiments. Bars show the mean ± S.D. with individual experiments overlaid. Mean values were 2.79% (control), 93.08% (GFP-CCDC81) and 25.83% (GFP-POC11). *n* = 3 independent experiments; *N* = 277 (control), 384 (GFP-CCDC81) and 347 (GFP-POC11) cells scored in total. P values were determined by one-way ANOVA with Dunnett’s post-hoc test against the untransfected control. **(E–H)** GFP-tagged CCDC81 truncations covering residues 1– 170 (E), 1–200 (F), 90–200 (G) and 200–800 (H). Transfected U2OS cells were stained for α-tubulin; GFP and α-tubulin are shown. 1–170 (E) and 1–200 (F) decorated microtubules, whereas 90–200 (G) and 200–800 (H) was diffusely cytoplasmic. The smallest fragment tested that retained microtubule binding was 1– 170. Scale bars= 5 µm. **(I)** Domain organization of CCDC81 (HU1, HU2 and the predicted coiled coil) with the truncations shown in (E–H) mapped beneath. Numbers are residues. **(J–L)** GFP-tagged POC11 truncations covering residues 251–450 (J), 300–450 (K) and 391–481 (L), treated and imaged as in (E–H). Only 391-481 (L) retained the microtubule binding. Scale bars = 5 µm. **(M)** Domain organization of POC11 with the truncations shown in (J–L) mapped beneath. Numbers are residues. **(N)** AlphaFold prediction of CCDC81 HU1–HU2 (residues 1-170) with an α/β-tubulin dimer. Left, predicted aligned error matrix; axes are residue number and the color scale is the expected error in the relative position of each residue pair, in Å. Right, the top-ranked model colored by chain (ipTM = 0.61 pTM = 0.73). The same model is shown in Figure 4Q; it is repeated here with its PAE matrix. **(O)** AlphaFold prediction of the CCDC81 C-terminal region (residues 747-800) with the POC11 N-terminal region in dimer (residues 1-51), shown as in (N) (ipTM = 0.59pTM = 0.65). The predicted disordered segment of CCDC81 is boxed in the domain diagram.

**Figure S7.**
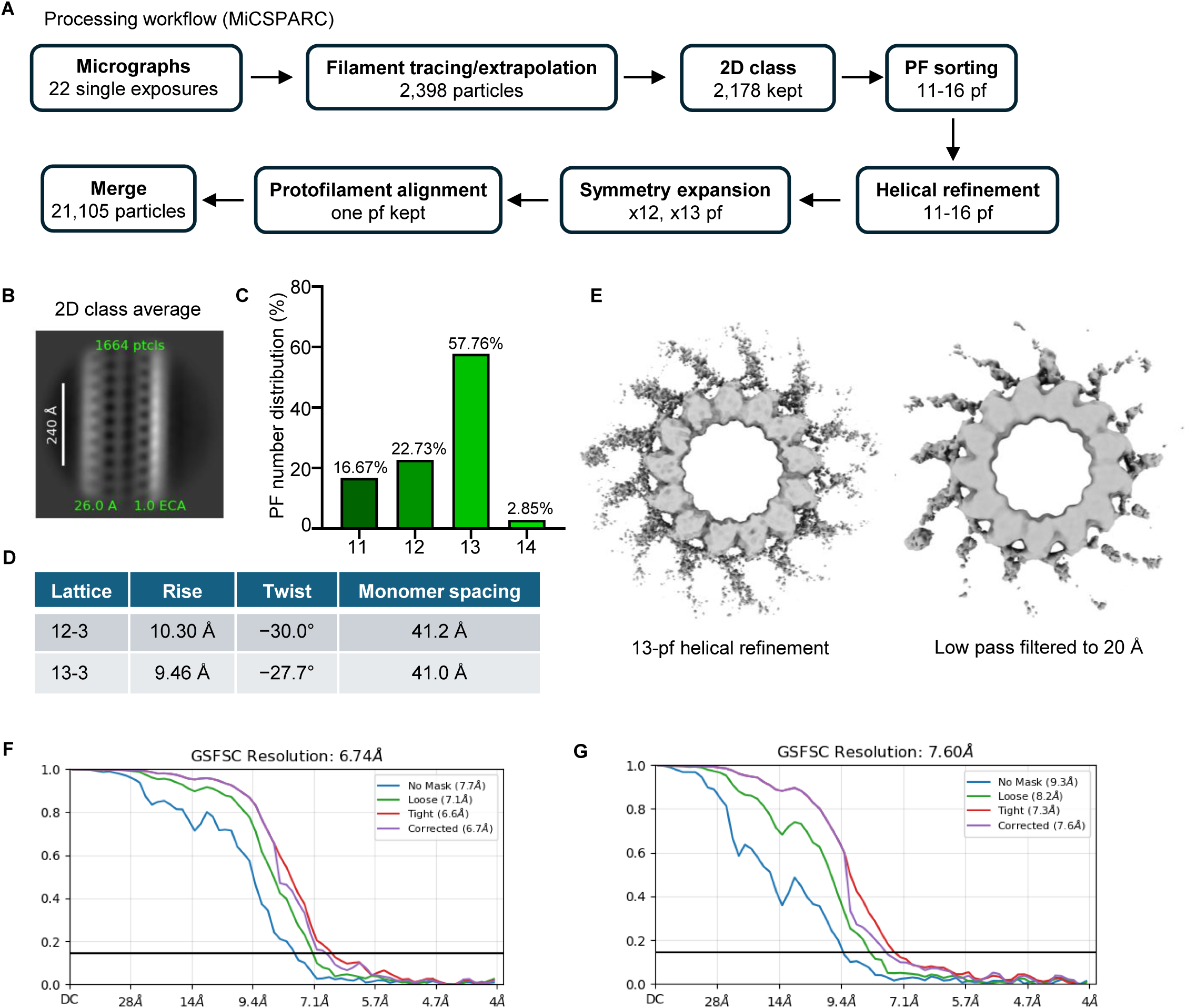
Cryo-EM processing of CCDC81-associated microtubules. **(A)** Processing workflow in CryoSPARC, following the MiCSPARC protocol. Numbers are the particles (microtubule segments) kept at each step. **(B)** Representative 2D class average (1,664 particles). Scale bar: 240 Å. **(C)** Distribution of protofilament numbers among the 2,178 particles after protofilament number sorting. **(D)** Helical parameters from the 12-3 and 13-3 helical refinements. Monomer spacing = rise x protofilament number ÷ helix start. **(E)** Top views of the 13-protofilament helical refinement, as refined (left) and low-pass filtered to 20 Å (right). **(F)** Gold-standard FSC curves for the map in Figure 5E. **(G)** Gold-standard FSC curves for class 0 shown in Figure 5F.

**Figure S8.**
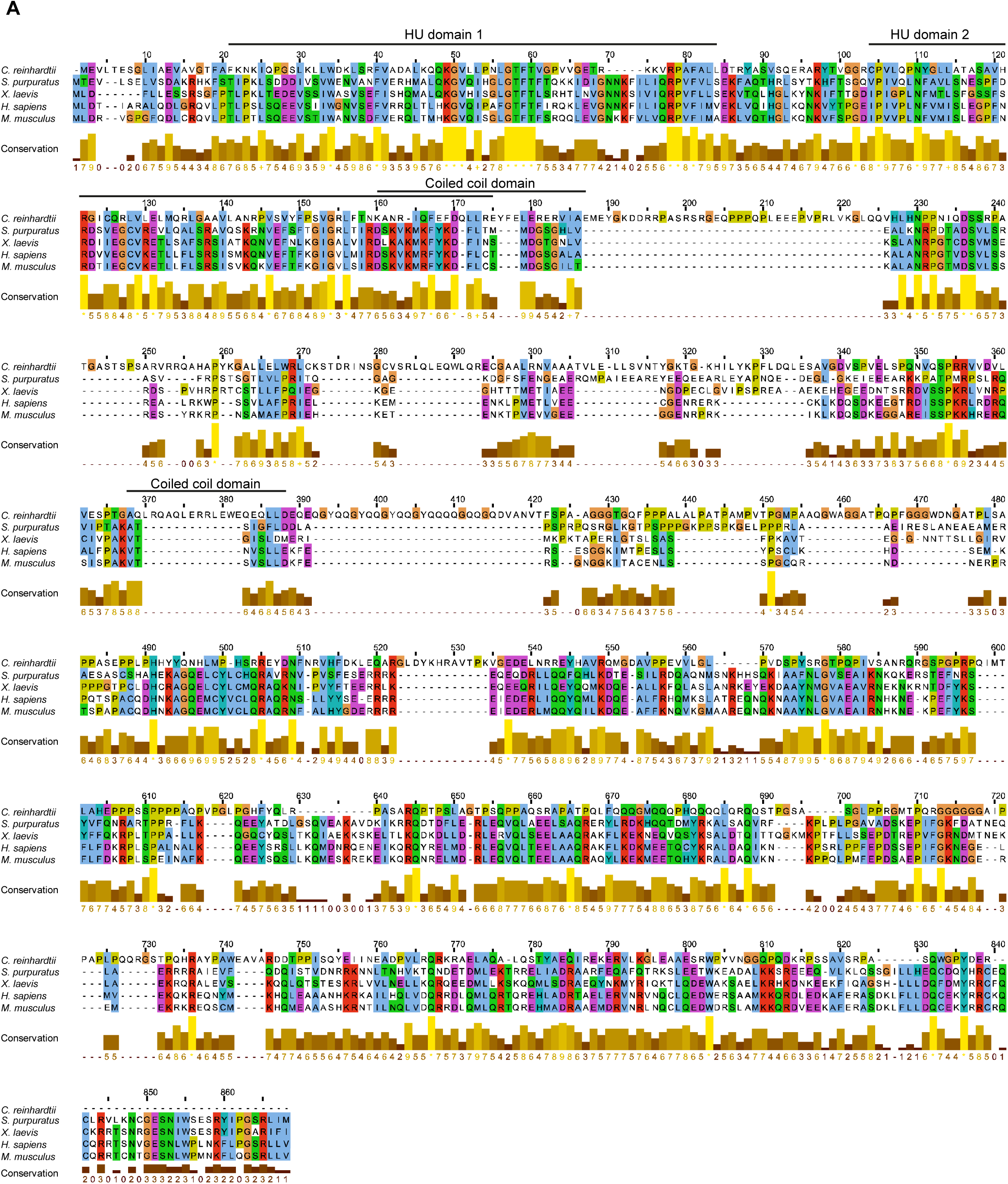
Sequence conservation of CCDC81 orthologues. **(A)** Alignment of CCDC81 sequences from *Chlamydomonas reinhardtii, Strongylocentrotus purpuratus*, *Xenopus laevis*, *Homo sapiens* and *Mus musculus*, generated with Clustal Omega and displayed in Jalview (version 2.7) and exported. Dashes indicate gaps and the numbering refers to alignment positions. The HU-1, HU-2 and predicted coiled-coil domains indicated above the alignment are those of CrCCDC81. The histogram beneath each block shows the Jalview conservation score for each alignment column; an asterisk marks columns that are identical in all five sequences and a plus sign marks columns in which all residues share the same physicochemical properties. Conservation scores were exported from Jalview and grouped into four categories to color the HsCCDC81 model shown in Figure 6B.

**Figure S9.**
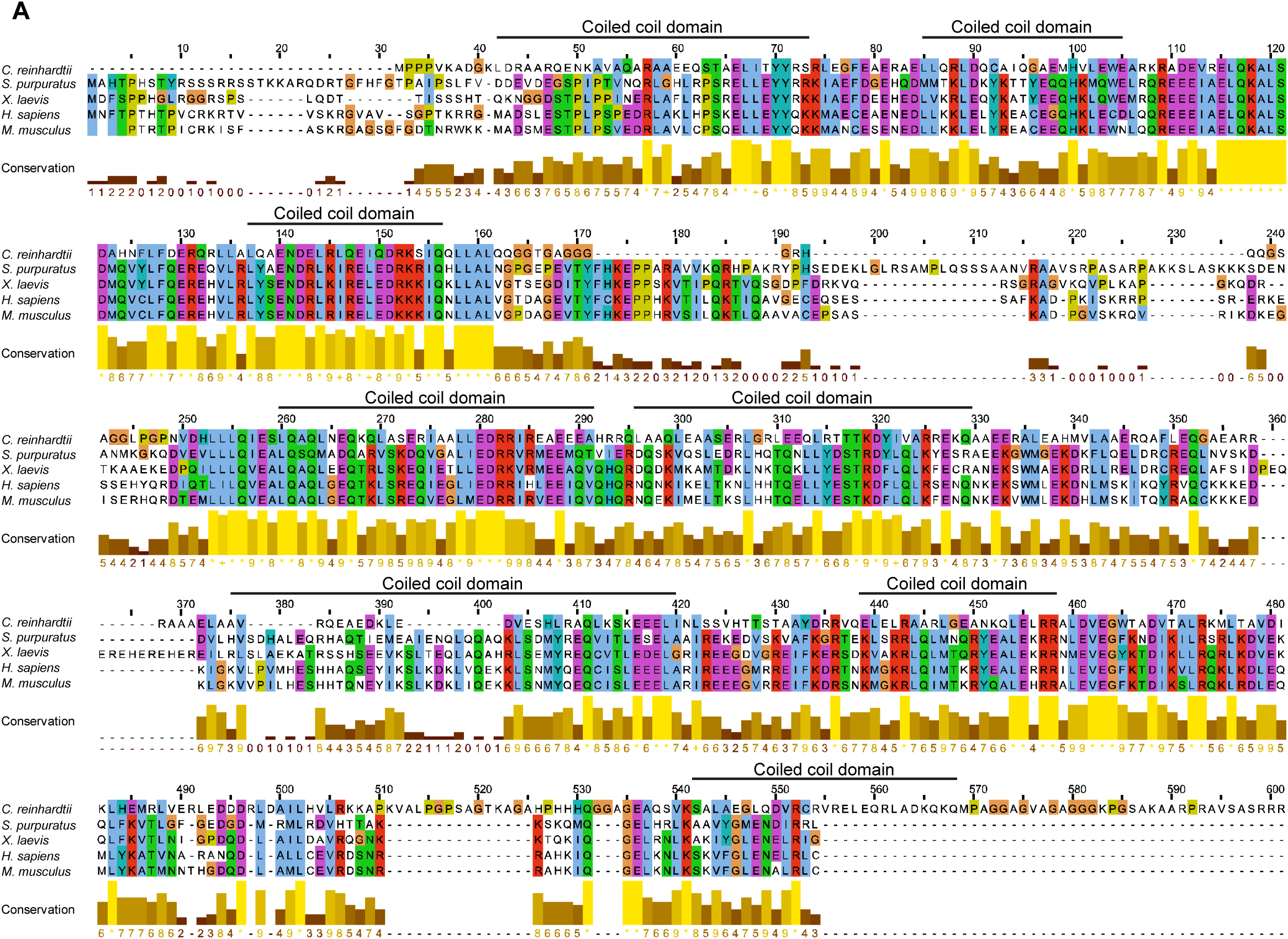
Sequence conservation of POC11 orthologues. **(A)** Alignment of POC11 sequences from *Chlamydomonas reinhardtii, Strongylocentrotus purpuratus*, *Xenopus laevis*, *Homo sapiens* and *Mus musculus*, generated with Clustal Omega and displayed in Jalview (version 2.7) and exported. Dashes indicate gaps and the numbering refers to alignment positions. The predicted coiled-coil domains indicated above the alignment are those of CrPOC11. The histogram beneath each block shows the Jalview conservation score for each alignment column; an asterisk marks columns that are identical in all five sequences and a plus sign marks columns in which all residues share the same physicochemical properties. Conservation scores were exported from Jalview and grouped into four categories to color the CCDC77 model shown in Figure S10F.

**Figure S10.**
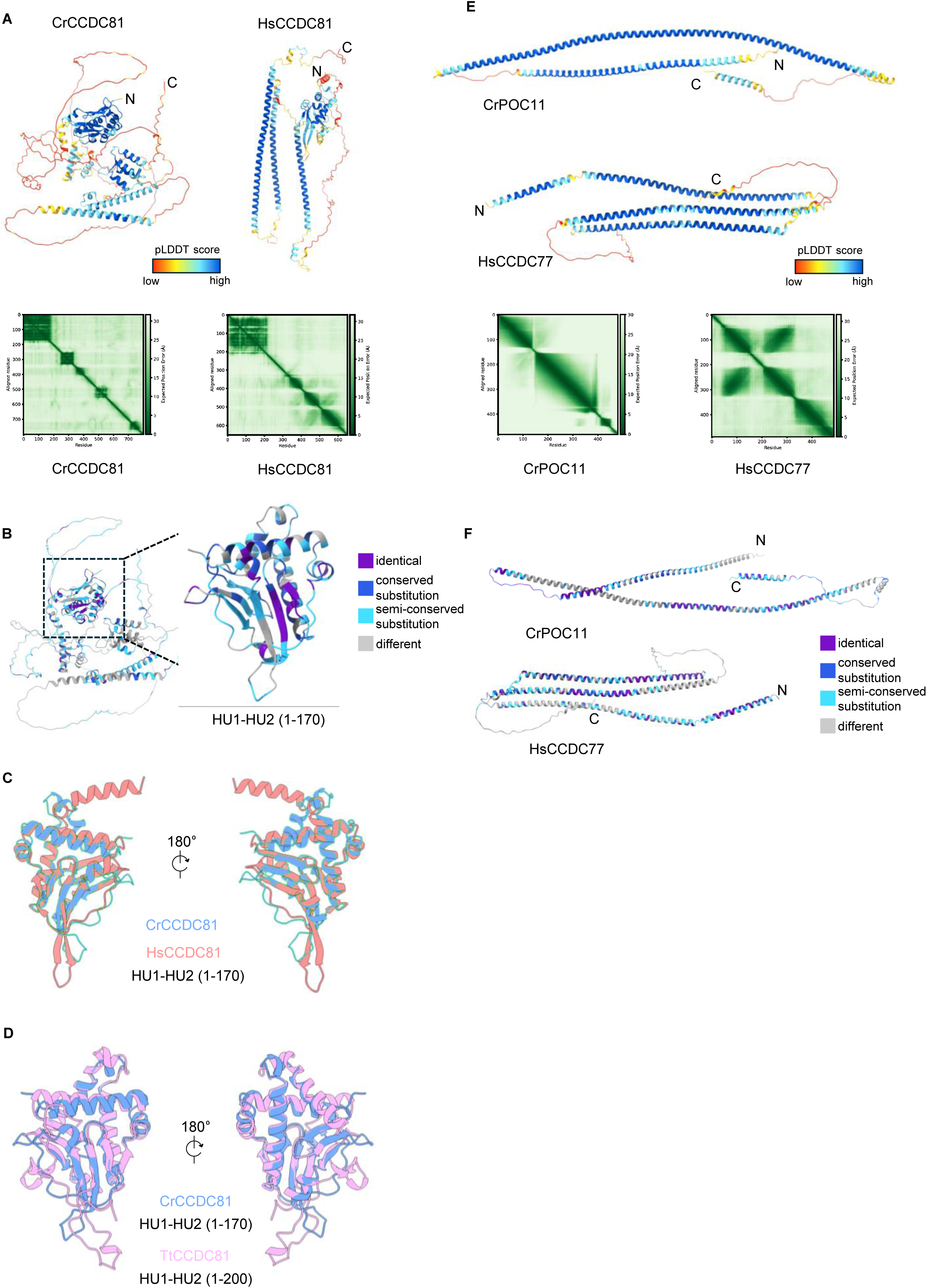
Structural predictions and comparisons of CCDC81 and POC11/CCDC77 orthologues. **(A)** AlphaFold models of CrCCDC81 and HsCCDC81 colored by pLDDT score, with the N and C termini indicated. Below, the predicted aligned error (PAE) matrices for the two models; axes are residue numbers and the color scale is the expected error in the relative position of each residue pair, in Å. **(B)** AlphaFold model of CrCCDC81 colored by sequence conservation, as in Figure 6B. The boxed region containing HU-1 and HU-2 (residues 1–170) is enlarged on the right. Conservation was scored from the alignment shown in Figure S8. **(C)** Superposition of the HU-1–HU-2 regions of CrCCDC81 (residues 1–170, blue) and HsCCDC81 (residues 1–170, salmon), shown in two views related by a 180° rotation. Models were superposed in ChimeraX using matchmaker with iterative pruning at a 2.0 Å cutoff. The structurally equivalent core superposes with a Cα RMSD of 1.11 Å over 100 residue pairs; across all 157 pairs, which include the divergent loops and termini, the RMSD is 6.73 Å. **(D)** Superposition of the HU-1–HU-2 regions of CrCCDC81 (residues 1–170, blue) and *Tetrahymena thermophila* CCDC81 (TtCCDC81, residues 1–200, pink), shown as in (C). The core superposes with a Cα RMSD of 1.09 Å over 85 residue pairs; across all 162 pairs the RMSD is 5.49 Å. The N-terminal region of TtCCDC81 binds microtubules. **(E)** AlphaFold models of CrPOC11 and HsCCDC77 colored by pLDDT score, with the N and C termini indicated. Below, the PAE matrices for the two models, shown as in (A). **(F)** AlphaFold models of CrPOC11 and HsCCDC77 colored by sequence conservation, scored from the alignment shown in Figure S9 as in (B). Conservation categories are as in Figure 6B.

**Figure S11.**
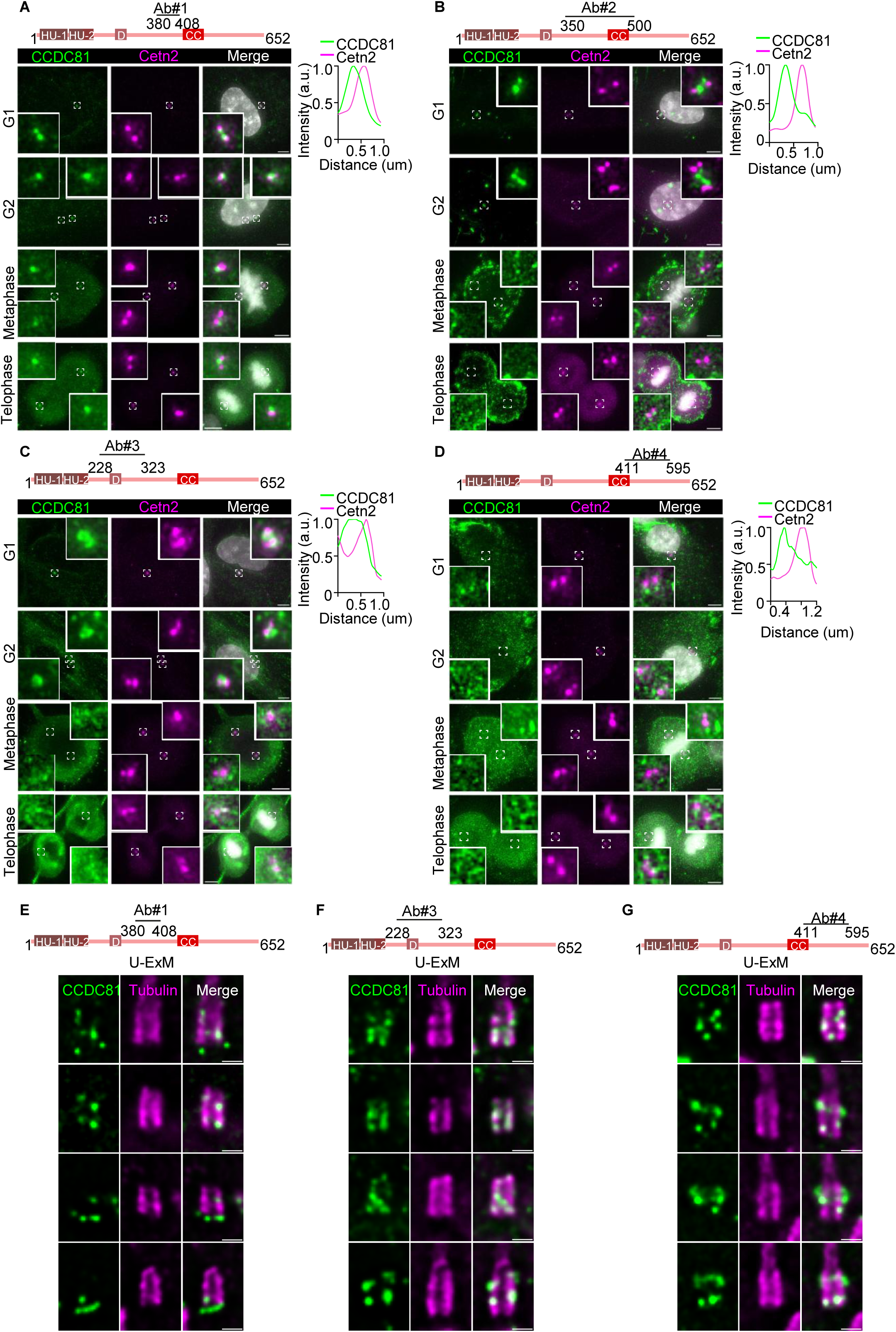
Evaluation of four independent antibodies against human CCDC81. **(A–D)** RPE-1 cells were fixed with methanol and stained for CCDC81 (green), Centrin-2 (Cetn2, magenta) and DNA (DAPI, grey), using Ab#1 (A; residues 380– 408), Ab#2 (B; residues 350–500), Ab#3 (C; residues 228–323) and Ab#4 (D; residues 411–595). Cell cycle stages were assigned from centriole number and DNA morphology. The region recognized by each antibody is mapped on the HsCCDC81 diagram above each panel. Insets are enlargements of the boxed regions at 4X. The plots on the right show intensity profiles for CCDC81 and Cetn2 measured across the centriole pair in the G1 inset, normalized to the maximum of each channel. Scale bars = 5 µm. **(E–G)** RPE-1 cells processed for U-ExM and stained for CCDC81 (green) and α/β tubulin (magenta), using Ab#1 (E), Ab#3 (F) and Ab#4 (G); each row shows an independent centriole. Ab#2 is shown in Figure 6C. Scale bars = 1 µm.

**Figure S12.**
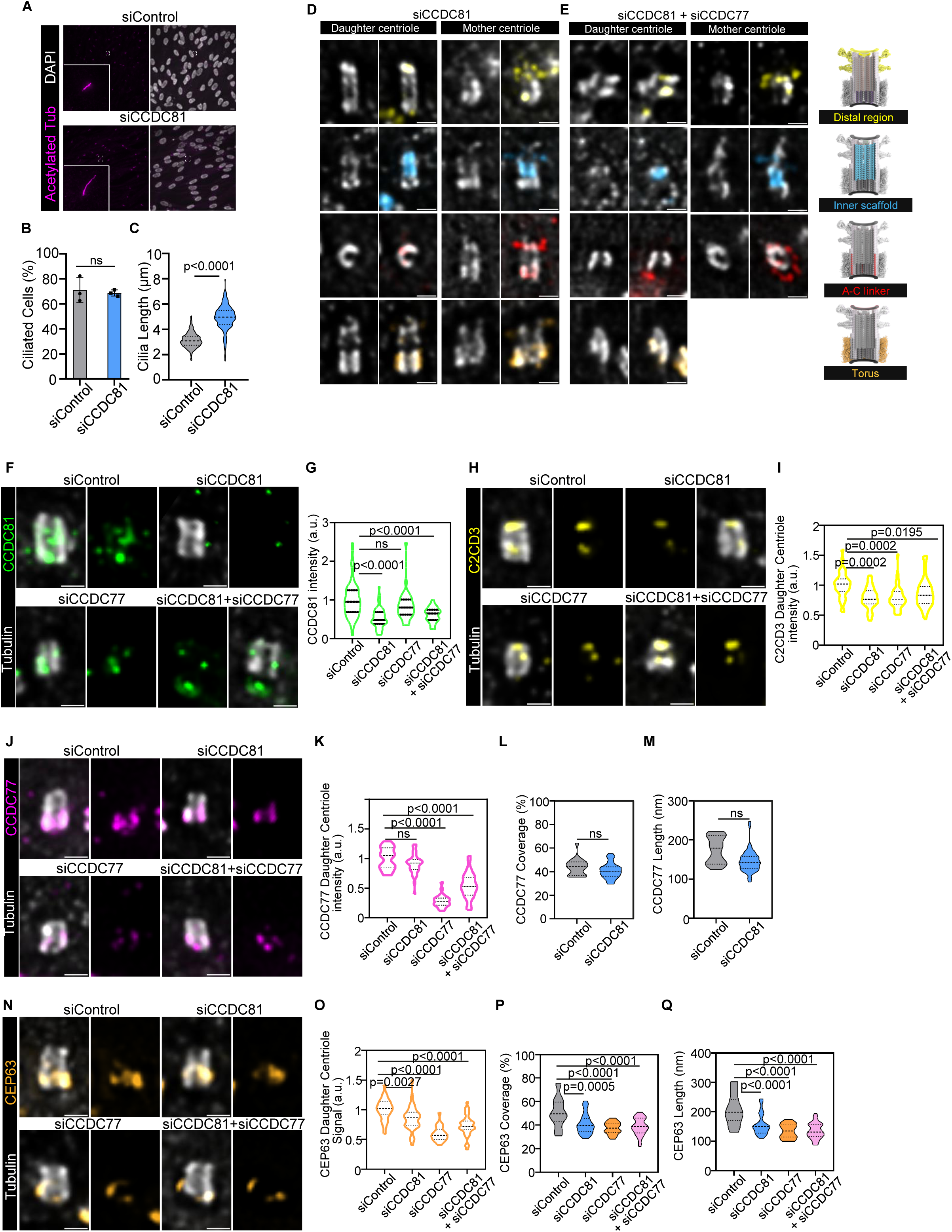
Ciliation, knockdown efficiency and additional centriolar markers in CCDC81 depleted cells. **(A)** siRNA transfected RPE-1 cells, serum starved and stained for acetylated tubulin (magenta) and DAPI (grey). Insets are enlargements of the boxed regions (4X). Scale bars = 5 µm. **(B)** Percentage of ciliated cells in each condition. Bars show the mean ± S.D. with individual experiments overlaid. siControl: N = 940, n = 3 (mean = 71.07%). siCCDC81: N = 1169, n = 3 (mean = 68.70%). **(C)** Cilia length measured from the acetylated tubulin signal. siControl: N = 357, n = 3 (mean = 3.135 µm). siCCDC81: N = 571, n = 3 (mean = 4.93 µm). **(D, E)** Additional examples of daughter and mother centrioles from siCCDC81 (D) and siCCDC81 + siCCDC77 (E) cells, processed for U-ExM and stained as in Figure 7A for α/β-tubulin (grey) together with C2CD3 (yellow), POC5 (cyan), WDR67 (red) or CEP63 (orange). Each centriole is shown as tubulin alone and as the merge. Scale bars = 1 µm. **(F)** Centrioles stained for α/β-tubulin (grey) and CCDC81 (green, Ab#2); for each condition, the merge and the CCDC81 channel alone are shown. Scale bars = 1 µm. **(G)** CCDC81 signal intensity on the daughter centriole, normalized to siControl. siControl: N = 107, n = 3 (mean = 1). siCCDC81: N = 140, n = 3 (mean = 0.53). siCCDC77: N = 98, n = 3 (mean = 0.84). siCCDC81 + siCCDC77: N = 64, n = 2 (mean = 0.61). **(H)** Daughter centrioles stained for α/β-tubulin (grey) and C2CD3 (yellow), shown as in (F). Scale bars = 1 µm. **(I)** C2CD3 signal intensity on the daughter centriole, normalized to siControl. siControl: N = 64, n = 3 (mean = 1). siCCDC81: N = 74, n = 3 (mean = 0.78). siCCDC77: N = 67, n = 3 (mean = 0.79). siCCDC81 + siCCDC77: N = 75, n = 3 (mean = 0.87). **(J)** Daughter centrioles stained for α/β-tubulin (grey) and CCDC77 (magenta), shown as in (F). Scale bars = 1 µm. **(K–M)** CCDC77 signal intensity on the daughter centriole, normalized to siControl (K), CCDC77 coverage (L) and the length of the CCDC77 signal along the centriole (M), defined as in Figure 7F–H. Coverage and length were measured only in siControl and siCCDC81 cells. (K) siControl: N = 36, n = 2 (mean = 1). siCCDC81: N = 70, n = 2 (mean = 0.91). siCCDC77: N = 47, n = 2 (mean = 0.28). siCCDC81 + siCCDC77: N = 55, n = 2 (mean = 0.54). (L) siControl: N = 34, n = 2 (mean = 43.82%). siCCDC81: N = 70, n = 2 (mean = 40.89%). (M) siControl: N = 34, n = 2 (mean = 174.0 nm). siCCDC81: N = 70, n = 2 (mean = 145.6 nm). **(N)** Daughter centrioles stained for α/β-tubulin (grey) and CEP63 (orange), shown as in (F). Scale bars = 1 µm. **(O–Q)** CEP63 signal intensity on the daughter centriole, normalized to siControl (O), the length of the CEP63 signal along the centriole (P) and CEP63 coverage (Q), defined as in Figure 6F–H. (O) siControl: N = 61, n = 3 (mean = 1). siCCDC81: N = 100, n = 3 (mean = 0.84). siCCDC77: N = 56, n = 3 (mean = 0.59). siCCDC81 + siCCDC77: N = 82, n = 3 (mean = 0.73). (P) siControl: N = 46, n = 3 (mean = 50.80%). siCCDC81: N = 82, n = 3 (mean = 41.06%). siCCDC77: N = 45, n = 3 (mean = 37.31%). siCCDC81 + siCCDC77: N = 77, n = 3 (mean = 38.61%). (Q) siControl: N = 46, n = 3 (mean = 207.2 nm). siCCDC81: N = 82, n = 3 (mean = 156.2 nm). siCCDC77: N = 45, n = 3 (mean = 134.5 nm). siCCDC81 + siCCDC77: N = 77, n = 3 (mean = 135.3 nm). Violin plots show the full distribution with the median and quartiles indicated. Lengths and scale bars in the U-ExM panels are corrected for the expansion factor (4X). *n* = number of independent experiments; *N* = number of cells (B), cilia (C) or centrioles (G, I, K–M, O–Q) per condition. P values were determined by one-way ANOVA with Dunnett’s post-hoc test against siControl, except in (L) and (M), where two-tailed t-test was used. ns, not significant.

**Table S1.**
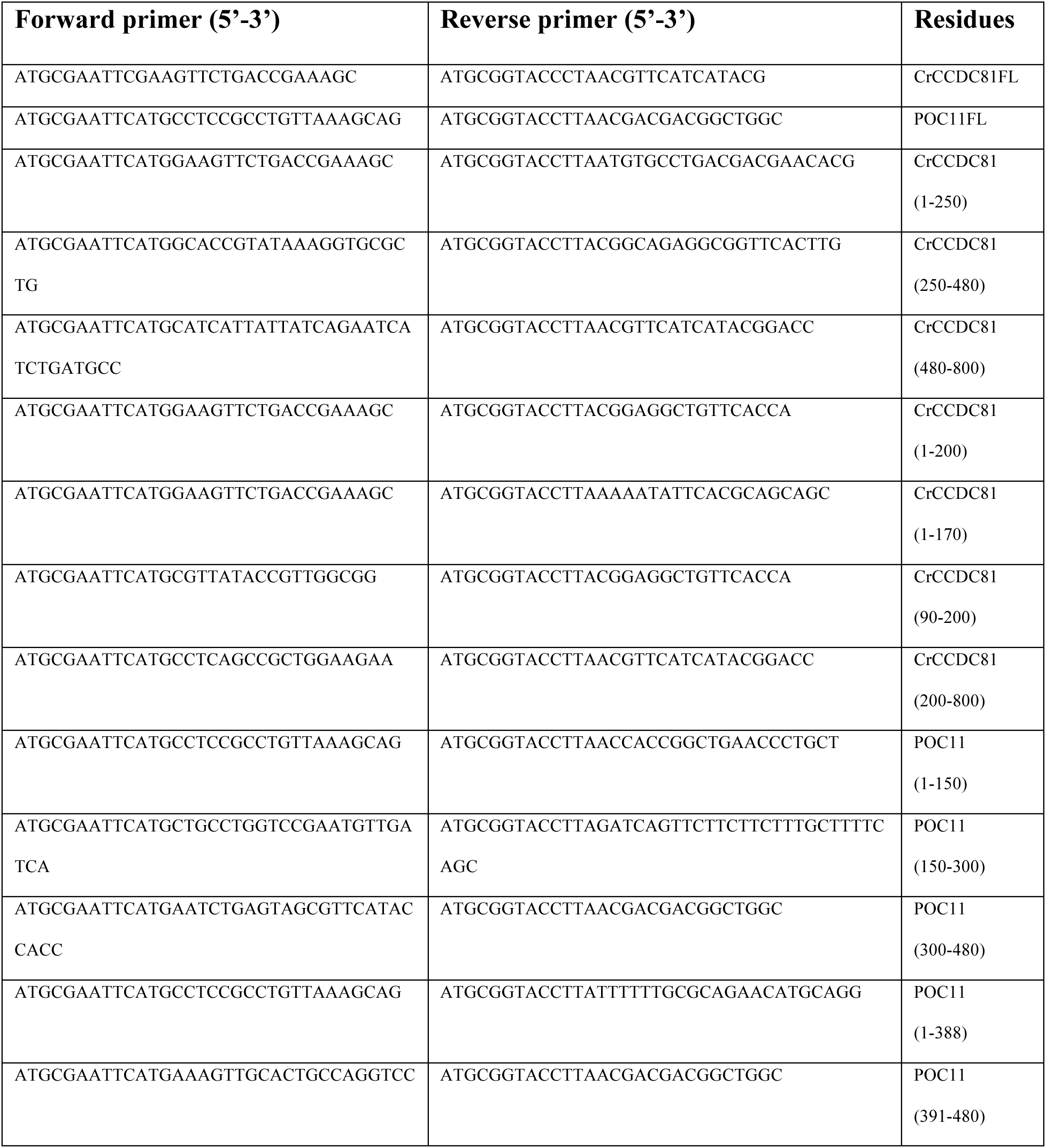
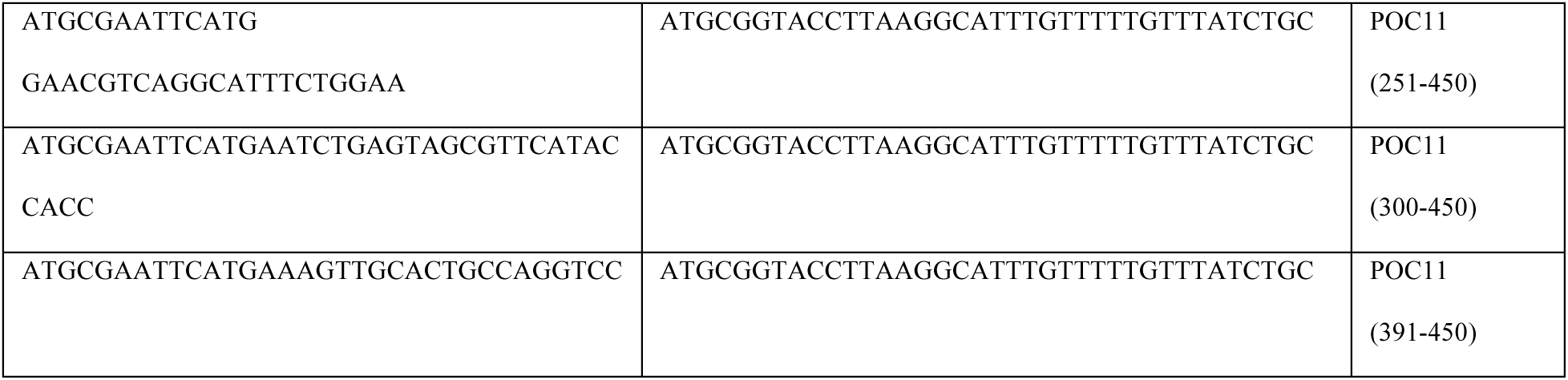
Primers used for CrCCDC81 and POC11 cloning into pEGFP-C2 and mCherry-C1 vectors.

**Table S2.** Uniprot accession numbers.

| **Gene** | **Species** | **Uniprot ID** |
|---|---|---|
| CCDC81 | *Homo sapiens* | Q6ZN84 |
| CCDC81 | *Chlamydomonas reinhardtii* | A0A2K3DWI7 |
| CCDC81 | *Mus musculus* | Q9D5W4 |
| CCDC81 | *Xenopus laevis* | A0A8J1MBB6 |
| CCDC81 | *Strongylocentrotus purpuratus* | A0A7M7PJJ3 |
| POC11 | *Homo sapiens* | Q9BR77 |
| POC11 | *Chlamydomonas reinhardtii* | A0A2K3DYN0 |
| POC11 | *Mus musculus* | Q9CZH8 |
| POC11 | *Xenopus laevis* | Q6DFC2 |
| POC11 | *Strongylocentrotus purpuratus* | A0A7M7N0C3 |

**Table S3.** Cryo-EM data collection and processing.

| **Data collection** | **Microtubules alone** | **Microtubules + MBP-CrCCDC81** |
|---|---|---|
| Microscope | Talos Arctica | Talos Arctica |
| Voltage (kV) | 200 | 200 |
| Detector | Falcon 4i, Selectris X energy filter (10 eV slit) | same |
| Magnification | 130,000x | 130,000x |
| Pixel size (Å) | 0.9024 | 0.9024 |
| Acquisition | Single exposures in linear mode, not dose-fractionated | same |
| Exposure time (s) | 9.2 | 9.2 |
| Electron exposure (e<sup>-</sup>/Å<sup>2</sup>) | 25-30 (mean 28) | 20-32 (mean 25) |
| Nominal defocus (µm) | -3.0 | -3.0 |
| Micrographs | 5 | 22 |

